# Ultrasound-reconfigurable scaffolds enable dynamic control of biochemical and biophysical cues for vascular network formation

**DOI:** 10.64898/2026.08.14.744961

**Authors:** Somnath Maji, Zubi Danish, Sam V. Varghese, Dharshan A. Hari, Abigail Pinch, Haijun Xiao, Carole Quesada, Andrew J. Putnam, Mario L. Fabiilli

## Abstract

Vascularization, which is critical for most engineered tissue constructs, is dependent on interactions between endothelial cells and spatiotemporally presented biochemical and biophysical cues. Yet, most hydrogels define these cues at the time of fabrication, thus precluding adjustments to actively drive vascular formation. We developed acoustically responsive scaffolds (ARSs) that use focused ultrasound to trigger growth factor release and localized matrix remodeling within fibrin hydrogels. ARSs were formed by incorporating phase-shift emulsions containing basic fibroblast growth factor (bFGF) along with perfluorohexane (C6) or perfluorooctane (C8). Upon ultrasound exposure, stable bubbles were generated in C6-ARSs that locally compacted the matrix and increased macroscale stiffness. Comparatively, in C8-ARSs, ultrasound generated macropores without impacting viscoelastic properties. Ultrasound increased bFGF release from both ARS types, which enhanced in vitro and in vivo vasculogenic assembly in ARSs with co-encapsulated endothelial cells and fibroblasts. Our data also show that ultrasound-driven matrix remodeling without bFGF release increased endothelial sprouting. In C6-ARSs, elevated levels of F-actin were observed in both cell types adjacent to bubbles as well as increased YAP intensity and nuclear asymmetry. Together, these results establish ARSs as reconfigurable hydrogels that couple on-demand release of biochemical cues with programmable matrix restructuring to direct three-dimensional microvascular assembly.

## 1. Introduction

Impaired or insufficient microvasculature is a central driver of morbidity in ischemic heart disease, stroke, and chronic limb ischemia; this leads to treatment complications and increased mortality [1–4]. Additionally, inadequate vasculature causes failure of many engineered tissue grafts, where lack of perfusion rapidly results in hypoxia, necrosis, and loss of function [5, 6]. This is especially problematic in thick or large volume constructs, which require rapid establishment of interconnected microvascular networks to survive and integrate after implantation, because diffusion sustains cell viability only within approximately 100-200 µm of a perfused vessel [7]. Pro□angiogenic growth factors and cell-based techniques have been evaluated to augment perfusion by stimulating vascular regeneration, but their translation has been hindered by short growth factor half-lives, poorly controlled dosing, and risks of aberrant or tumor-like neovascularization [8, 9]. Collectively, these limitations highlight the need for new approaches that actively orchestrate microvascular network formation in three dimensions rather than relying solely on passive host responses [5].

Tissue engineering has utilized vasculogenesis, the de novo assembly of endothelial cells into blood vessels, to generate microvascular networks within biomaterial scaffolds. Hydrogels containing fibrin, collagen, gelatin methacryloyl, and polyethylene glycol have emerged as leading platforms because they mimic key features of the extracellular matrix and support 3D endothelial network formation, particularly when combined with appropriate stromal cells [10–14]. Collectively, these systems show that vasculogenesis is exquisitely sensitive to biochemical cues - such as hypoxia, gradients of vascular endothelial growth factor (VEGF) and basic fibroblast growth factor (bFGF), and paracrine signaling - as well as biophysical cues including matrix stiffness, viscoelasticity, plasticity, and architectural confinement [10, 15–17]. For example, stiffness and viscoelasticity regulated tip-cell specification and sprouting via yes-associated protein (YAP)-dependent mechanotransduction, while temporal changes in matrix mechanics and hypoxia sequentially modulated endothelial cluster-based vasculogenesis [15, 18, 19]. Despite these advances, most hydrogel-based, vasculogenic platforms impart or embed relevant cues only at the time of fabrication, thus limiting the ability to dynamically adjust growth factor release profiles or microstructural features after fabrication and/or implantation.

Stimuli-responsive “smart” biomaterials offer a promising route to overcome this bottleneck, particularly when coupled to external triggers that are already clinically integrated. Focused ultrasound is especially attractive as a noninvasive stimulus because it can penetrate deeply within the body, can be focused with millimeter precision, and is routinely used in diagnostic and therapeutic procedures [20]. Building on these advantages, we have developed composite hydrogels termed acoustically responsive scaffolds (ARSs) that consist of a hydrogel (e.g., fibrin) doped with phase□shift emulsion. The micron-sized emulsion droplets contain a perfluorocarbon liquid that undergoes a phase-shift into a vapor when exposed to ultrasound above a threshold amplitude. This non-thermal, phase-shift is termed acoustic droplet vaporization (ADV) [21]. In our prior studies, ADV was used to release growth factors, which were initially encapsulated within the emulsion, into the matrix. Focused ultrasound enables spatiotemporally controlled release of growth factors like bFGF from an ARS, which stimulates spatially directed angiogenesis [22] and restoration of perfusion in the murine hind-limb ischemia model [23]. Sequential payload release from ARSs has further demonstrated that ultrasound can gate multiple factors (e.g., bFGF followed by platelet derived growth factor) with distinct pressure and/or frequency thresholds, underscoring the tunability of this platform [24, 25]. In addition to drug release, ADV also locally restructures the hydrogel matrix, with generated features dependent on the perfluorocarbon liquid within the emulsion. For example, emulsions containing perfluorohexane (C_6_F_14_) form stable bubbles that persist within the hydrogel matrix following ultrasound exposure. Within strain-stiffening hydrogels (e.g., fibrin, collagen), the bubbles locally compact and stiffen the matrix [26]. Alternatively, bubbles generated from emulsions containing perfluorooctane (C_8_F_18_) recondense following ultrasound exposure. This produces macropores that are accessible to liquid, micron-sized particles, and cells [27]. Prior ARS studies have largely focused on acellular scaffolds and on bulk measures of angiogenesis, with limited interrogation of how ADV in a cell-laden, fibrin matrix reorganizes the microarchitecture, modulates 3D vasculogenesis, or engages mechanotransduction pathways in the context of vasculogenic co-cultures. Thus, up until now, it remained unknown whether ARSs could be leveraged not simply as growth-factor depots, but as actively reconfigurable microenvironments that coordinate biochemical release and mechanical remodeling to control vasculogenic assembly in three dimensions.

In this study, we investigated the effects of ADV-mediated release of bFGF, a potent inducer of blood vessel formation, and local matrix restructuring on vasculogenic assembly in a fibrin-based ARS. We hypothesized that by varying the perfluorocarbon species within the phase-shift emulsion and ultrasound exposure history, ADV would create spatially heterogeneous, architectural niches within the matrix that modulate endothelial behavior and activity; concurrently, ADV-triggered release of bFGF would provide a coordinated biochemical cue. To test this, we investigated the effect of two perfluorocarbon species on bFGF release kinetics from an ARS. We characterized ADV-induced matrix restructuring using confocal microscopy and bulk rheology. Additionally, we characterized the biocompatibility and vasculogenic self-assembly of ARSs containing a co-culture of endothelial cells and fibroblasts. Studies were conducted in vitro and in vivo in a subcutaneous mouse model. By integrating mechanobiological readouts with structural and functional measures of vasculogenesis, this work aims to establish ARSs as a new class of bioactive materials capable of actively shaping microvascular network formation through coupled biochemical and biophysical modulation of the 3D microenvironment.

## 2. Materials and Methods

### Cell Culture

Human umbilical vein endothelial cells (HUVECs, C2519A, Lonza, Walkersville, MD, USA) and HUVECs expressing red fluorescent protein (RFP-HUVECs, cAP-0001RFP, Angio-Proteomie, Boston, MA, USA) were cultured in fully supplemented endothelial growth media (VascuLife VEGF Endothelial Medium Complete Kit, Lifeline Cell Technology, Frederick, MD, USA). Normal human dermal fibroblasts (NHDFs, Lonza) were cultured in Dulbecco’s modified Eagle medium (DMEM, Gibco, Carlsbad, CA, USA) supplemented with 10% (v/v) fetal bovine serum (Corning, Glendale, AZ, USA), 100 U/mL penicillin, and 100 μg/mL streptomycin (Thermo Fisher Scientific, Waltham, MA, USA). Cells were grown to 70% confluency before subculturing. All experiments in this study were carried out with passage numbers 3-4 for both HUVECs and RPF-HUVECs as well as 8-9 for NHDFs.

### Microfluidic preparation of phase-shift double emulsion

Double emulsions with a water-in-perfluorocarbon-in-water (W_1_/PFC/W_2_) structure were produced by a microfluidic technique as previously described [36]. Perfluorohexane (C6, C_6_F_14_, CAS# 355-42-0, Strem Chemicals, Newburyport, MA, USA) or perfluorooctane (C8, C_8_F_18_, CAS# 307-34-6, Sigma-Aldrich, Burlington, MA USA) was the PFC phase. Fluorosurfactant copolymer, derived from a 2:1 molar ratio of Krytox 157 FSH (CAS# 51798-33-5, DuPont, Wilmington, DE, USA) and poly(ethylene glycol)bis(amine) (MW: 1000 g/mol, CAS# 24991-53-5, Alfa Aesar, Ward Hill, MA, USA), was dissolved in PFC at 2% (w/w) by vortexing. The PFC solution was combined at 2:1 (v/v) with a W_1_ solution containing 0.15□mg/mL bFGF (100-18B, Thermo Fisher Scientific) and 1 mg/mL bovine serum albumin in 5 mM Tris buffer (pH 7.6). In some studies, bFGF was omitted from the W_1_ phase. The PFC and W_1_ phases were sonicated (Q55 with CL-188 immersion probe, QSonica, LLC, Newton, CT, USA) for 30□s while on ice to produce the primary emulsion (W_1_/PFC). Double emulsions were generated by flowing the primary emulsion and W_2_ phase, 50 mg/mL Pluronic F68 in phosphate buffered saline (PBS) through a glass microfluidic chip (3200136, junction: 14 μm × 17 μm, Dolomite, Royston, United Kingdom) at 1 μL/min and 10 μL/min, respectively (**Fig. 1A**). Characterization using a Coulter Counter (Multisizer 4, Beckman Coulter, Brea, CA, USA) equipped with a 50 μm aperture tube revealed that the double emulsions had an average diameter, coefficient of variation, and concentration of 7.3 ± 0.4 μm, 26.8 ± 2.8%, and (2.1 ± 0.2) × 10^9^ particles per mL, respectively.

**Fig. 1.**
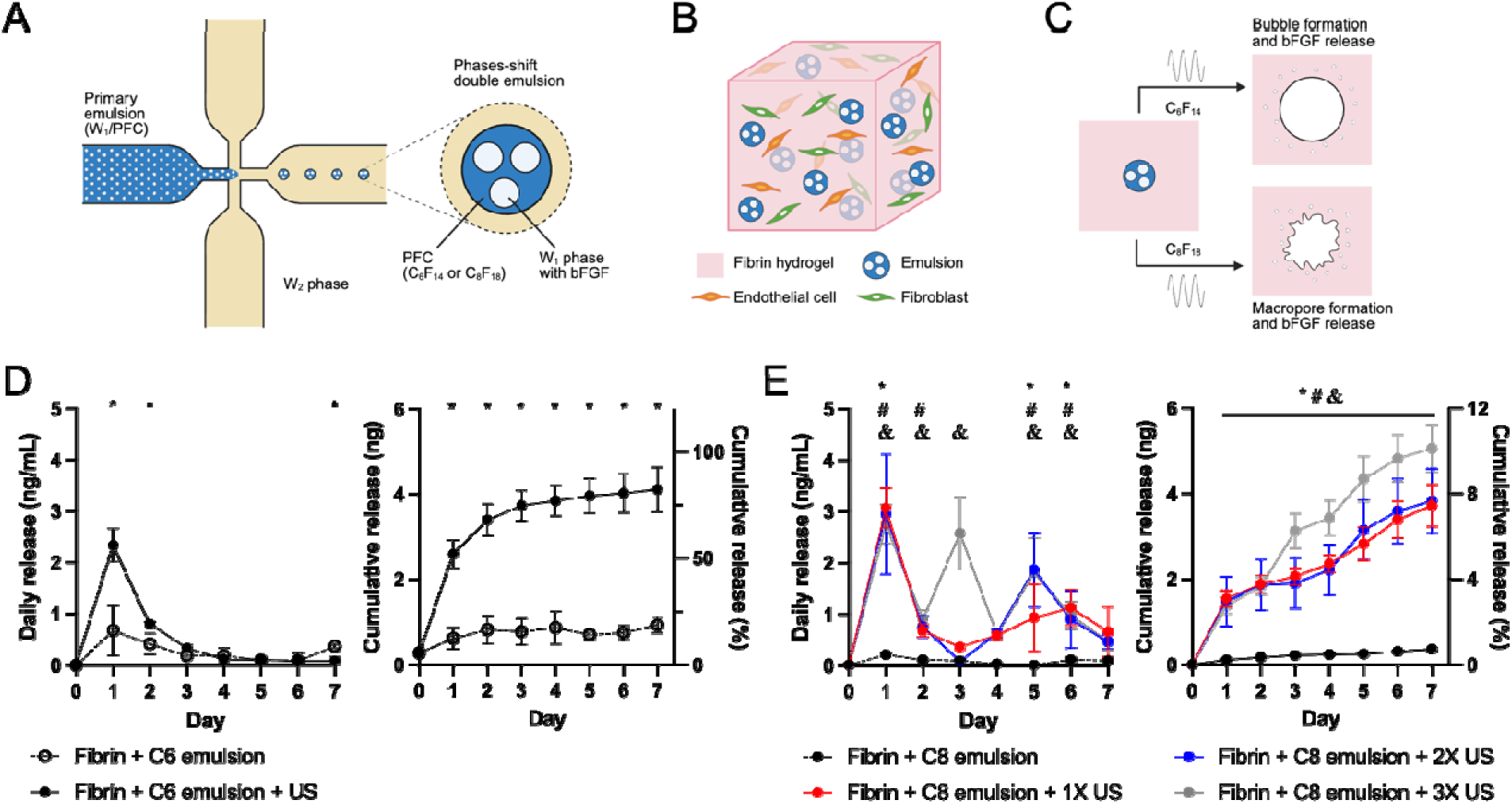
The phase-shift double emulsion within an ARS enables ultrasound-mediated drug release and matrix restructuring. (A) Schematic depicting the microfluidic generation of phase-shift double emulsions. bFGF was dissolved within the W_1_ phase of the primary emulsion (W_1_/PFC), which was subsequently encapsulated by the W_2_ phase to form a double emulsion. (B) Schematic of a cell-loaded ARS that contained both endothelial cells and fibroblasts. (C) A schematic shows that ultrasound (US) exposure yielded bubbles or macropores in the ARS when the PFC phase consisted of C_6_F_14_ or C_8_F_18_, respectively. (D,E) For both PFCs, bFGF wa released by US. Daily and cumulative bFGF release profiles for acellular ARSs containing emulsions with C_6_F_14_ (D) or C_8_F_18_ (E). In panel D, US was applied on day 0. In panel E, US wa applied on day 0 (+1X US), days 0 and 4 (+2X US), or days 0, 2, and 4 (+3X US). N = 4 gels per group. Significant differences are denoted as follows: no US vs. +US (*, panel D); no US vs. +1X US (*, panel E), no US vs. +2XUS (#, panel E), and no US vs. +3X US (&, panel E).

### Fabrication of ARSs

Bovine fibrinogen (F8630, Sigma-Aldrich) was dissolved in DMEM at 20 mg/mL clottable protein while under gentle vortex mixing for 30 seconds. The fibrinogen solution and additional DMEM were degassed separately in a vacuum chamber at ∼6 kPa for 60 minutes to minimize the amount of dissolved gas, which could act as cavitation nuclei. ARSs were made by combining the prepared fibrinogen, DMEM, 0.02% (v/v) C6 emulsion or 0.2% (v/v) C8 emulsion, and human recombinant thrombin (Recothrom, Baxter, Deerfield, IL, USA). The final concentrations of fibrinogen and thrombin in the ARSs were 5 mg/mL and 2 U/mL, respectively. To prepare cell-loaded ARSs (**Fig. 1B**), HUVECs or RFP-HUVECs and NHDFs were trypsinized and mixed with the fibrin solution at the desired concentration (specified below). To enable visualization of the matrix, 75 μg/mL Alexa Fluor 647-labeled fibrinogen (Invitrogen, Waltham, MA, USA) was added to each ARS prior to polymerization. Acellular or cell-loaded, fibrin-only gels (i.e., without emulsion) were prepared similarly. Acellular gels (0.5 mL volume) were polymerized in a 24-well BioFlex plate (Flexcell International, Burlington, NC, USA). Cell-loaded ARSs (0.2 mL) were prepared in a custom plastic plate (well diameter: 9.5 mm, well height: 5.5 mm) sealed with a Tegaderm membrane (3M Healthcare, St. Paul, MN, USA). Gels polymerized for 10 min at room temperature and were subsequently covered with either 0.5 mL media (BioFlex plate) or 0.2 mL media (custom plate). Following exposure to ultrasound, gels in custom plates were transferred to a standard 24-well plate and covered with 0.5 mL media. Unless otherwise noted, gels were stored in a tissue culture incubator at 37O°C with 5% carbon dioxide throughout the experiments.

### Ultrasound exposure setup and parameters

Ultrasound experiments were conducted in a tank (32 cm × 61 cm × 32 cm) filled with degassed, deionized water at 37 °C. ADV was induced using a calibrated, focused transducer (2.5 MHz, H-108, f-number: 0.83, radius of curvature: 50 mm, Sonic Concepts, Bothell, WA, USA). Pulsed waveforms (peak negative pressure: 4.2 MPa, pulse duration: 12 μs; pulse repetition frequency: 100 Hz) were generated by a function generator (33500B, Agilent Technologies, Santa Clara, CA, USA) and amplified by a gated radiofrequency amplifier (GA-2500A, Ritec, Warwick, RI, USA). The amplified signals were monitored in real-time on an oscilloscope (HDO4034, Teledyne LeCroy, Chestnut Ridge, NY, USA).

In vitro experiments were performed by submerging the custom plate containing ARSs in the water tank; alternatively, the BioFlex plate was placed in the tank such that only the bottom contacted the water. The transducer, which was placed beneath the plate, was connected to a three-axis positioning system controlled by MATLAB (The MathWorks, Natick, MA, USA). To generate ADV, the transducer was rastered at 3 mm/s with 0.5 mm spacing between raster lines. The raster pattern was completed on three planes by placing the axial focus from +3 mm to +1 mm above the well bottom.

### Measurement of bFGF release

Acellular ARSs were polymerized in BioFlex plates, covered with 0.5 mL media consisting of DMEM with 100 U/mL penicillin and 100 μg/mL streptomycin, and exposed to ultrasound using the aforementioned exposure conditions. ARSs were exposed to ultrasound either one (day 0), two (days 0 and 4), or three (days 0, 2, and 4) times. The overlying media were sampled daily for 7□days. The concentration of bFGF in the collected samples was measured using an enzyme-linked immunosorbent assay (DY233, R&D System, Inc., Minneapolis, MN, USA).

### ARS morphology and mechanical properties

Acellular ARSs containing Alexa Fluor 647 fibrin were imaged with a confocal microscope (AX, Nikon, Melville, NY, USA). For ARSs with C6 emulsion (**Fig. 1C**), bubble diameter and the width of the fibrin matrix compacted by the bubble were measured in NIS-Elements (Nikon). For ARSs with C8 emulsion, the following metrics were evaluated: volume fraction of macropores within the ARS, major/minor axis of each macropore, and macropore density. Storage and loss moduli of ARSs were characterized using a rheometer (HR 10, TA Instruments, New Castle, DE, USA) with a parallel plate geometry (measurement head diameter: 8 mm). Using an axial force of 0.3 N, ARSs were interrogated at 1 Hz oscillatory shear and 1% strain rate, which was previously determined to be in the linear viscoelastic regime [28].

### Cellular viability and metabolic activity assays

To assess cellular viability, 2×10^5^ RFP-HUVECs per mL and 2×10^5^ NHDFs per mL were encapsulated within ARSs and cultured in fully supplemented endothelial growth media. Constructs were stained with 167 nM Sytox green (Invitrogen) in Hank’s balanced Salt solution (Gibco) for 20 min, followed by counterstaining with Hoechst 33342 (Invitrogen). Constructs were then imaged with confocal microscopy. Using a custom script (NIS Elements), the number of Hoechst+, RFP+ and Sytox+ cells was calculated to quantify total cells, HUVECs, and dead cells within each gel. The number of NHDFs was determined based on the difference between Hoechst+ and RFP+ cells.

Metabolic activity of the constructs was assessed using the alamar blue assay (Invitrogen) as done previously[20]. Constructs were incubated in 10% (v/v) alamar blue reagent in DMEM for 4 h. After incubation, fluorescence of the supernatants was measured at 560 nm and 590 nm using a plate reader (SpectraMax M2e, Molecular Devices, San Jose, CA, USA) and analyzed according to the manufacturer’s instructions. The metabolic activity of ARSs containing a monoculture of 2×10^5^ NHDFs per mL, with different concentrations of emulsions, was also investigated.

### In vitro vasculogenesis studies

HUVECs and NHDFs were encapsulated within fibrin-only gels or ARSs at 5 x 10^5^ cells per mL for each cell type. A dose response study was conducted to characterize the effect of bFGF concentration on vasculogenesis. bFGF was incorporated into starvation media consisting of DMEM/F12 supplemented with 10□µg/mL insulin, 5□µg/mL transferrin, 6.7□ng/mL sodium selenite, 100□U/mL penicillin, 100□µg/mL streptomycin, 100□µg/ml ovalbumin (Sigma-Aldrich), 50□µg/ml l-ascorbic acid (Sigma-Aldrich), 1□µg/ml hydrocortisone 21-hemisuccinate sodium salt (Sigma-Aldrich), and 0.75□U/mL heparin. Cell-loaded, fibrin-only gels were cultured for 7 days in either starvation media, starvation media containing bFGF, or fully supplemented VascuLife media.

To assess ADV-mediated release of bFGF on vasculogenesis, ARSs were formulated with HUVECs, NHDFs, and bFGF-loaded emulsion. Constructs were cultured in starvation media for 7 days. To isolate the effect of ADV-induced matrix restructuring, cell-loaded ARSs with 10^6^ cells per mL for each cell type were fabricated with blank emulsion (i.e., without bFGF) and cultured in fully supplemented VascuLife media. For both studies, ARSs were exposed to ultrasound on day 0 (i.e., the same day as polymerization).

### In vivo studies

All animal procedures were approved by the University of Michigan Institutional Animal Care and Use Committee. Male immunocompromised SCID hairless mice (SHO strain; N = 15, 4–6 weeks old, 22.7 ± 1.2 g, Charles River Laboratories, Wilmington, MA, USA) were utilized as hosts. Mice were anesthetized with isoflurane and the dorsal skin was scrubbed with povidone-iodine. Two full-thickness longitudinal incisions were made in the lower dorsal region on either side of the spine, creating subcutaneous pockets. Cell-loaded ARSs or fibrin-only gels (volume = 0.25 mL) were implanted into the pockets, and the surgical sites were closed with sutures. A subset of constructs was treated with ultrasound on the day of implantation. During treatment, each mouse was anesthetized with isoflurane and placed in a supine position on an exposure platform positioned on the water tank. Construct morphology was monitored using B-mode ultrasound on a ZS3 system (Mindray, Mahwah, NJ, USA) with a 24 MHz linear array (L30-8, Mindray). Construct height and average pixel intensity were measured in Fiji. The implanted constructs and surrounding tissue were harvested on day 7 post implantation.

### Cell staining and analyses

Constructs were fixed in 4% (w/v) paraformaldehyde. In vitro constructs were stained with rhodamine labelled Ulex europaeus agglutinin I (UEA-I, 1:100, RL-1062-2, Vector Laboratories, Newark, CA, USA) overnight at 4°C. Constructs were then permeabilized with 0.1% (v/v) Triton X-100 and incubated overnight with either Alexa Fluor 488-labeled phalloidin (1:400, Invitrogen) or primary antibody against VE-cadherin (1:400, ab318152, Abcam, Cambridge, United Kingdom). An Alexa Fluor 488-labeled secondary antibody (1:1000, ab150061, Abcam) was used for visualization in addition to Hoechst 33342 for counterstaining nuclei.

In vivo samples were embedded in paraffin, sectioned, and mounted onto glass slides by the University of Michigan Orthopaedic Research Laboratories Histology Core. Histological slides were stained with hematoxylin and eosin (H&E). The H&E-stained slides were used to measure the diameter of bubbles and macropore sizes in Fiji. In vivo constructs were stained for UEA-I and immunohistochemically stained for YAP (1:2000, ab205270, abcam), Ter-119 (1:200, 14-5921-82, Invitrogen), CD31(1:1600, ab182981, abcam), and nucleolin (1:100, 4691-RBM4, NeoBiotechnologies, Union City, CA, USA). Sections were subsequently counterstained with 4′,6-diamidino-2-phenylindole (DAPI, 1:500).

Stained samples were imaged with a confocal microscope and custom scripts (NIS Elements) were used to quantify marker expression. For in vitro samples, the following metrics were calculated: total length of UEA-I+ tubules per volume of hydrogel (i.e., tubule density, mm/mm^3^), length of each UEA-I+ tubule versus distance from nearest bubble edge, summed F-actin intensity per volume of HUVEC or NHDF, and summed VE-cadherin intensity per length. For in vivo samples, the following metrics were calculated: %UEA-I area (i.e., UEA+ area normalized by implant area), % Ter-119 area (i.e., Ter-119+ area normalized by implant area), % CD31 area (i.e., CD31+ area normalized by granulation tissue area), nuclear circularity versus distance from nearest bubble edge, and average YAP intensity in cell versus distance from nearest bubble edge. Additionally, the density of host cells that migrated into each implant was calculated based on the difference between the number of DAPI+ and nucleolin+ cells.

### Statistical analysis

Statistical analyses were performed using GraphPad Prism software (GraphPad Software, Inc., La Jolla, CA, USA). For experiments comparing only two groups, we used two-tailed, unpaired Student’s t-tests. For experiments comparing multiple groups, we used one-way ANOVA and Tukey’s multiple comparisons test. Unless otherwise noted, data are presented as mean ± standard deviation.

## 3. Results

### 3.1. Ultrasound increased bFGF release from fibrin ARSs

We first characterized bFGF release from ARSs containing either C6 or C8 emulsions under no ultrasound or ultrasound exposure conditions (**Fig. 1D, 1E**). In the absence of ultrasound, C6-ARSs released a low amount of bFGF over 7 days, with a peak on day 1 followed by near-baseline release thereafter. In contrast, exposure to a single ultrasound treatment on day 0 caused a marked increase in release on day 1, followed by a rapid decline over subsequent days. Cumulative release reached approximately 4.1 ng by day 7, compared with approximately 0.8 ng in the corresponding no-ultrasound control (**Fig. 1D**).

A similar ultrasound-dependent increase in bFGF release was observed in C8-ARSs, but release kinetics differed from C6-ARSs. Without ultrasound, minimal daily release was seen across the 7-day period, remaining near baseline. A single ultrasound exposure on day 0 (i.e., 1X US) produced a pronounced increase on day 1, followed by sustained release through day 7. Repeated ultrasound exposures on day 4 (i.e., 2X US) as well as on days 2 and 4 (i.e., 3X US) further altered the release profile, with additional release peaks after subsequent insonation events. By day 7, cumulative release was significantly higher in the 3X ultrasound group compared to 1X and 2X groups (**Fig. 1E**). Repeated exposures were not conducted for the C6-ARSs since high densities of ultrasound-generated bubbles interfere with the propagation of ultrasound in subsequent insonation events, thus providing no additional release [29].

Comparison of the two emulsions demonstrated that ultrasound enhanced bFGF release in both ARS formulations, but the temporal profiles were formulation dependent. The C6 emulsion produced a predominantly early, transient release response after ultrasound, whereas the C8 emulsion supported a more sustained and restimulatable release pattern across multiple insonation events. Together, these data show that both ultrasound exposure and emulsion chemistry contribute to controlling bFGF release from fibrin ARSs, establishing the basis for subsequent cell-based analyses.

### 3.2. Ultrasound tuned matrix architecture non-invasively

Next, we characterized ADV-induced features within C6-ARSs. Before ultrasound exposure, the emulsion appeared as discrete droplets within the fibrin network (**Fig. 2A, 2B**). After ultrasound exposure, the droplets were converted into stable bubbles that radially compacted the surrounding matrix and elevated the fluorescence signal within the fibrin; bubbles persisted through day 7. Bubbles increased significantly in size over time, as seen qualitatively in confocal images and quantitatively (**Fig. 2C**). In parallel, the width of the compacted, fibrin rim surrounding the bubbles also increased over time, with no significant difference between day 1 and day 4, but a significant increase by day 7 (**Fig. 2D**). These images and measurements indicate progressive expansion of the bubble-containing regions within the gel.

**Fig. 2.**
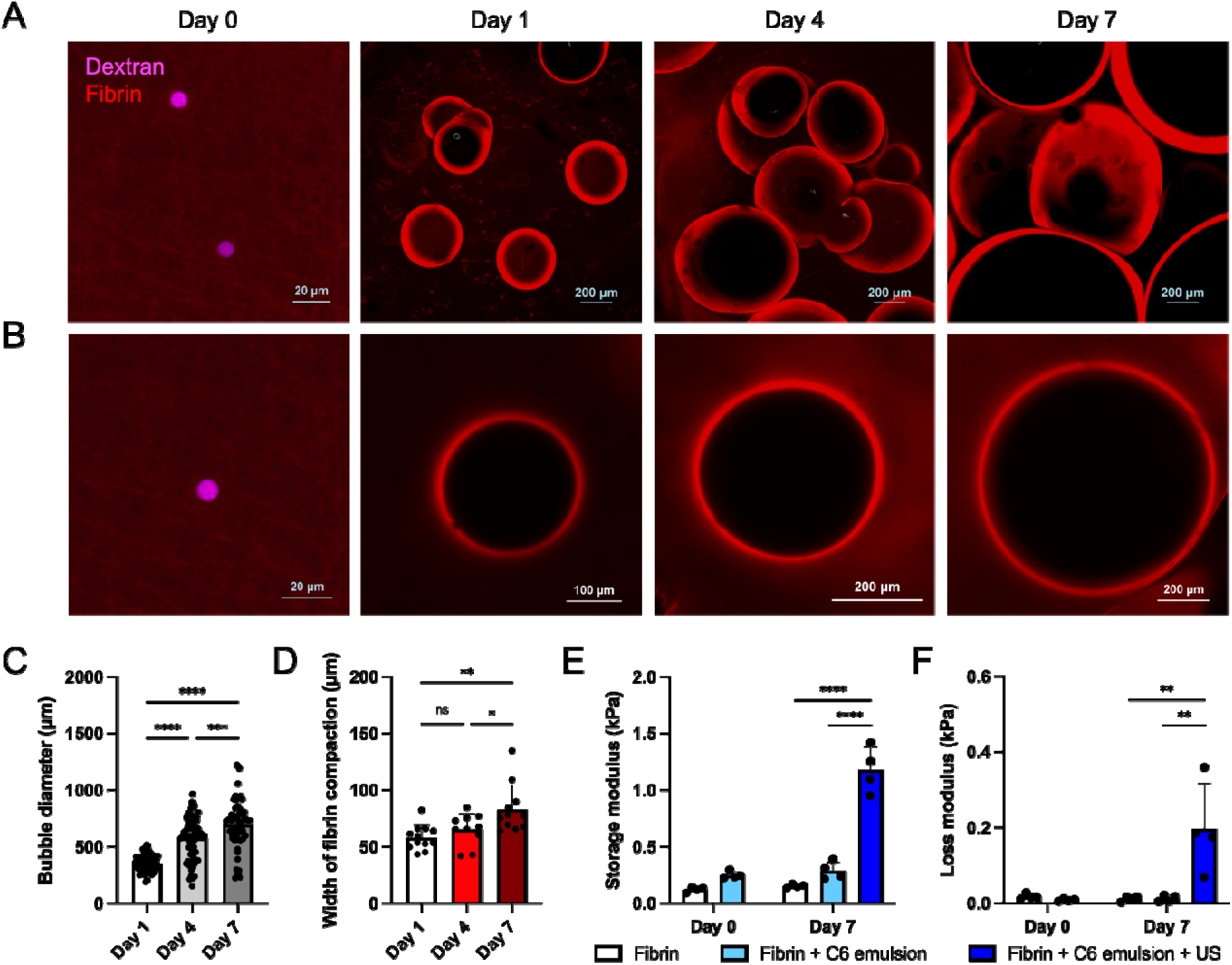
Ultrasound generated stable bubbles within C6-ARSs that locally compact the fibrin matrix and cause macroscale stiffening. (A) Maximum intensity projections and (B) single-plane confocal images show longitudinal bubble growth after ultrasound (US) exposure. Day 0 images reveal the presence of double emulsion droplets, containing a fluorescent dextran payload in the W_1_ phase, prior to ultrasound (US) exposure. (C) Bubble diameter and (D) width of the compacted fibrin rim surrounding each bubble were quantified based on confocal images. N = 4 gels per group with 4-5 fields of view per gel. (E) Storage and (F) loss moduli increased after ultrasound exposure and bubble formation. N=4 gels per group. Significant differences are denoted as follows: *p<0.05, **p<0.01, ***p<0.001, and ****p<0.0001; ns denotes a non-significant difference.

Rheological analysis showed that ultrasound-treated C6-ARSs became mechanically stiffer after exposure, with the storage modulus increasing significantly on day 7 relative to day 0 and corresponding non-ultrasound controls (**Fig. 2E**). The loss modulus also increased after ultrasound, although it remained lower than the storage modulus (**Fig. 2F**). In contrast, fibrin-only gels and C6-ARSs without ultrasound showed non-significant changes over the same period. Together, these data highlight that in C6-ARSs, ultrasound induced formation of stable bubbles that grew over time and concurrently compacted the fibrin matrix, leading to an increased scaffold stiffness.

We also evaluated ADV-induced features within C8-ARSs (**Fig. 3A, 3B**). In the absence of ultrasound, the fibrin gel remained largely uniform; ultrasound generated macropores throughout the scaffold that appeared more irregular in shape compared to bubbles within C6-ARSs, though they also had a bright periphery. Given the ability to repeatedly stimulate ADV at multiple timepoints (**Fig 1E**), we evaluated how repeated ultrasound exposure impacted macropore formation. Macropore volume correlated with the number of ultrasound exposures, with the group receiving ultrasound on days 0, 2, and 4 (i.e., 3X US) exhibiting the largest macropore volume (**Fig. 3C**). Pore density decreased with the number of ultrasound exposures (**Fig. 3D**), highlighting consolidation of the formed macropores. In parallel, both the major (**Fig. 3E**) and minor (**Fig. 3F**) axes of the macropores increased with the number of exposures, indicating enlargement in multiple dimensions. Together, these measurements indicate progressive remodeling of the scaffold architecture with repeated acoustic activation.

**Fig. 3.**
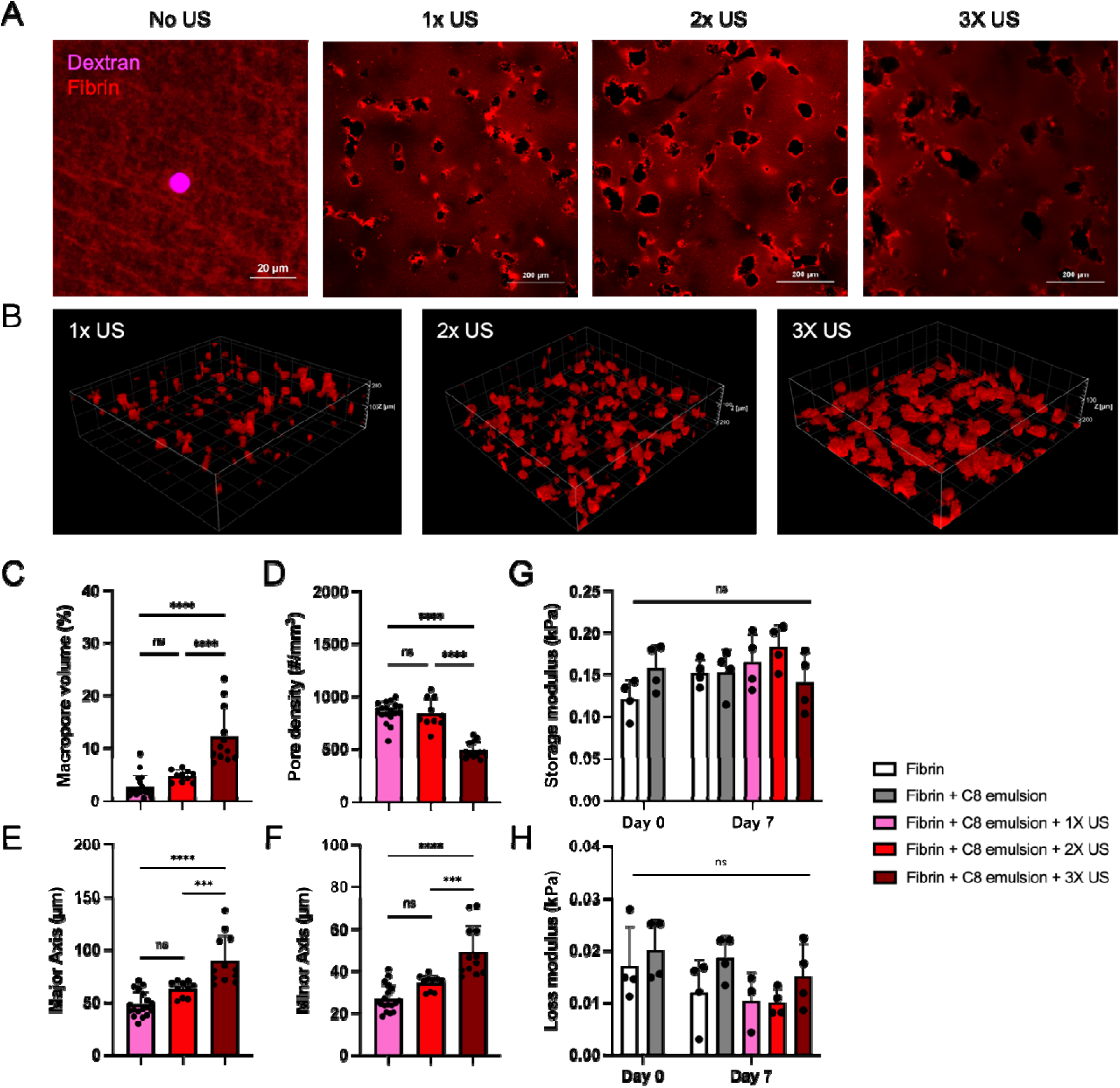
Ultrasound generated macropores within C8-ARSs without altering bulk viscoelasticity. (A) Single-plane confocal images and (B) 3D reconstructions of z-stacks show macropores following varying numbers of ultrasound (US) exposures. In panel A, the fibrin matrix is shown in red; an emulsion droplet containing fluorescent dextran in the W_1_ phase i observed in the no US condition. In panel B, macropores are denoted in red. Macropore volume (C) and density (D) correlated directly and inversely with the number of US exposures, respectively. The major (E) and minor (F) axes of the macropores also correlated with the number of US exposures. N = 4 gels per group with 4-5 fields of view per gel. Rheological analysis (N=4 gels per group) showed no significant differences in storage (G) or loss (H) moduli. Significant differences are denoted as follows: ***p<0.001 and ****p<0.0001; ns denotes a non-significant difference.

Despite these marked changes in macropore morphology, rheological testing showed no significant difference in either storage (**Fig. 3G**) or loss (**Fig. 3H**) modulus across the tested conditions. Thus, ultrasound-induced macropore formation altered the microarchitecture of the C8-ARSs without significantly changing bulk viscoelastic properties.

### 3.3. ADV was cytocompatible for cells within ARSs

To identify a suitable concentration of emulsion, we evaluated the metabolic activity of NHDFs encapsulated within C6- or C8-ARSs with different emulsion concentrations (**Fig. S1**). Metabolic activity increased longitudinally for all groups during the 7-day study. There were no significant differences when comparing matched sonicated versus non-sonicated groups at any time point. On day 7, sonicated C6-ARSs with 0.01% and 0.02% (v/v) emulsion as well as C8-ARSs with 0.2% (v/v) emulsion exhibited the highest metabolic activities. As such, 0.02% and 0.2% (v/v) were selected as the emulsion concentrations for all subsequent studies.

We quantified the viability of RFP-HUVECs and NHDFs co-encapsulated within ARSs to assess the cytocompatibility of ADV-mediated remodeling (**Fig. 4, S2**). Confocal images show abundant Hoechst+ nuclei and RFP-HUVECs across all conditions, with some dead cells (i.e., Sytox+) at each time point. In general, fibroblasts displayed greater viability than RFP-HUVECs on days 1 and 4. There was a longitudinal increase in viability for both cell types, with RFP-HUVECs exhibiting greater gains. For both fibrin-only gels and C6-ARSs, ultrasound exposure did not significantly alter viability compared to the corresponding non-ultrasound controls. On day 7, the viability of NHDFs and RFP-HUVECs was 99% and 85%, respectively, in the sonicated C6-ARS group. For C8-ARSs, ultrasound exposure did not impact the viability of either cell type on days 1 and 4. On day 7, there still was no significant difference between unexposed C8-ARSs and those that received a single ultrasound exposure. However, there was a significant difference for C8-ARSs that received two and three exposures, with the latter yielding viabilities of 79% and 78% for NHDFs and RFP-HUVECs, respectively. We also evaluated the cellular metabolic activity of the co-culture-laden gels (**Fig. S3**). Relative fluorescence units increased over time from day 1 to day 7 across all conditions, with no major loss of metabolic activity after ultrasound exposure. Overall, these data indicate that ADV-mediated remodeling of fibrin ARSs is compatible with cell survival and metabolic activity in both monoculture and co-culture settings, supporting its use for subsequent vasculogenic experiments.

**Fig. 4.**
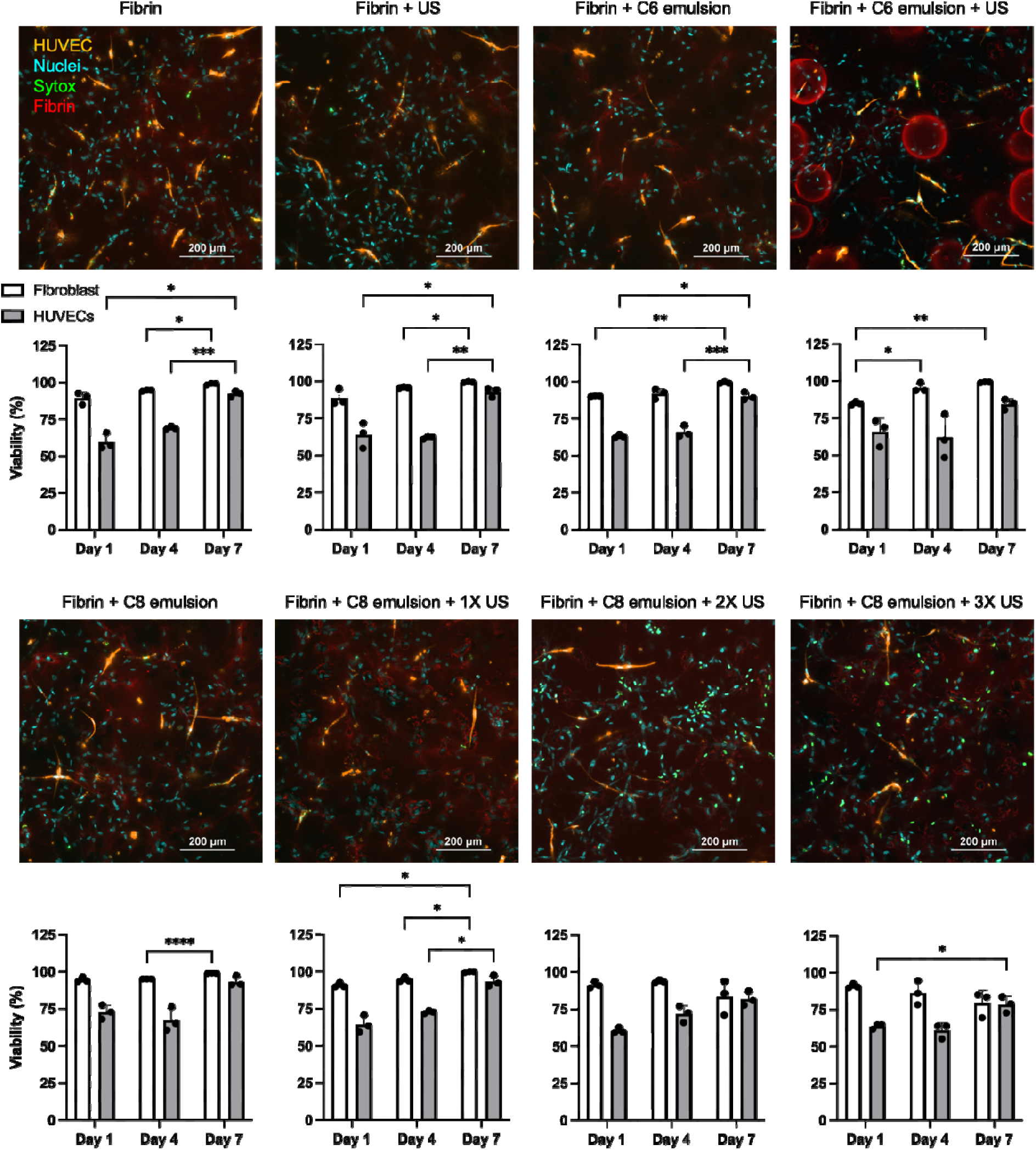
Cell viability was preserved following ultrasound exposure in ARSs. Representative confocal images show RFP-HUVECs and NHDFs co-encapsulated within fibrin-only gels as well as ARSs with C6 or C8 emulsion after 7 days of culture. Viability was assessed using Sytox-staining. Below each image is the longitudinal viability of each cell type for each experimental condition. N=3 gels per group with 5-10 fields of view per gel. Significant differences ar denoted as follows: *p<0.05, **p<0.01, ***p<0.001, and ****p<0.0001; ns denotes a non-significant difference. Additional images and statistical analyses are found in Fig. S2.

### 3.4. ADV-triggered bFGF release promoted vasculogenic assembly in vitro

Prior to evaluating the effect of ADV-mediated bFGF release on endothelial sprouting and assembly, we performed a bFGF dose response study in fibrin-only gels containing HUVECs and NHDFs (**Fig. 5A, S4**). There was a correlation between tubule density and bFGF concentration, with 1-100 ng/mL yielding densities significantly greater than the negative control (i.e., starvation media without bFGF) though lower than the level obtained in fully supplemented media. Thus, vasculogenic assembly was stimulated by the presence of bFGF within the media.

**Fig. 5.**
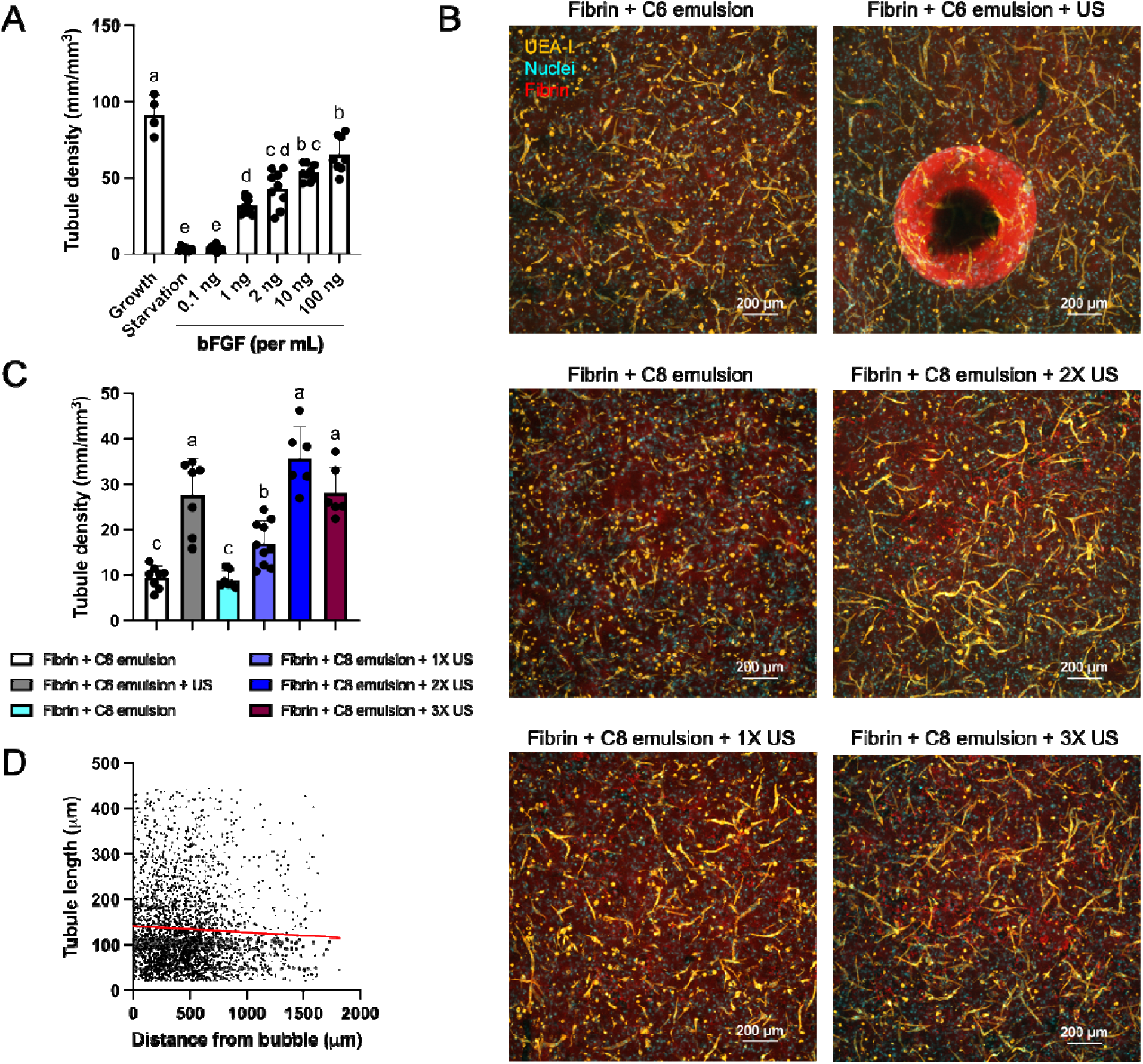
Ultrasound-triggered bFGF release in ARSs enhances endothelial sprouting. (A) Prior to ultrasound (US) studies, HUVECs and NHDFs were co-encapsulated in fibrin-only gel and cultured for 7 days in either growth media, starvation media, or starvation media supplemented with varying concentrations of bFGF. Tubule density was assessed via quantification of UEA-I staining. Examples images are found in Fig. S4. N =4-9 gels per group with 5-10 fields of view per gel. (B) Representative confocal images show UEA-I-stained endothelial structures in ARSs, which contained bFGF-loaded emulsions, after 7 days of culture in starvation media (C). US exposure increased tubule density in both C6- and C8-ARSs. N=6-10 gels per group with 5-10 fields of view per gel. (D) Following US exposure in C6-ARSs, tubule length correlated inversely with distance from the bubble interface. The linear trend, shown by the red line, is significantly negative (p = 0.006). N=3555 endothelial sprouts from N=7 gels. Statistically significant differences in panels A and C are denoted in compact letter display.

Next, we co-cultured HUVECs and NHDFs within ARSs for 7 days after ultrasound exposure. In these studies, bFGF was encapsulated within the phase-shift emulsion and ARSs were cultured in starvation media. Representative confocal images showed denser and more organized endothelial networks in the ultrasound-treated groups than in the corresponding non-ultrasound controls (**Fig. 5B**). Based on a quantitative analysis of the UEA-I stained images, there were significant increases in tubule density when comparing ultrasound-treated versus untreated groups for both C6- and C8-ARSs (**Fig. 5C**). With C8-ARSs, tubule density increased going from one to two ultrasound exposures; however, there was no significant difference when comparing two versus three exposures. Consequently, all subsequent studies utilized two ultrasound exposures for C8-ARSs. We also investigated whether tubule length exhibited a spatial dependence relative to the ADV-induced features within C6-ARSs. Quantification of tubule length as a function of distance from the bubble interface showed a modest though significantly negative linear trend (p = 0.006), with longer tubules observed nearer the bubble (**Fig. 5D**). Together, these data show that ultrasound-triggered bFGF release in fibrin ARSs increased endothelial tubule assembly, with repeated insonation further modulating the response in C8-ARSs.

### 3.5. Tubule density was enhanced by ADV-generated bubbles and macropores

To test whether matrix remodeling contributed to endothelial responses, we formulated ARSs with blank emulsion (i.e., without bFGF), HUVECs, and NHDFs and then cultured the constructs for 7 days in fully supplemented media. Confocal imaging revealed that ultrasound-treated C6- and C8-ARSs formed significantly greater endothelial networks compared to the fibrin-only gel without ultrasound exposure (**Fig. 6A, 6B, S5**). In the sonicated C6-ARS group, the intensity of F-actin (**Fig. 6A**) increased significantly for both NHDFs (**Fig. 6C**) and HUVECs (**Fig. 6D**); no significant difference was observed for the sonicated C8-ARS group. In contrast, VE-cadherin levels were comparable across all groups, with no significant differences detected (**Fig. 6E, 6F**). Together, these data show that ultrasound-driven remodeling of the fibrin matrix increased endothelial network formation even under full growth conditions. These results support the interpretation that an ARS influences endothelial organization through structural and mechanobiological cues in addition to growth-factor release.

**Fig. 6.**
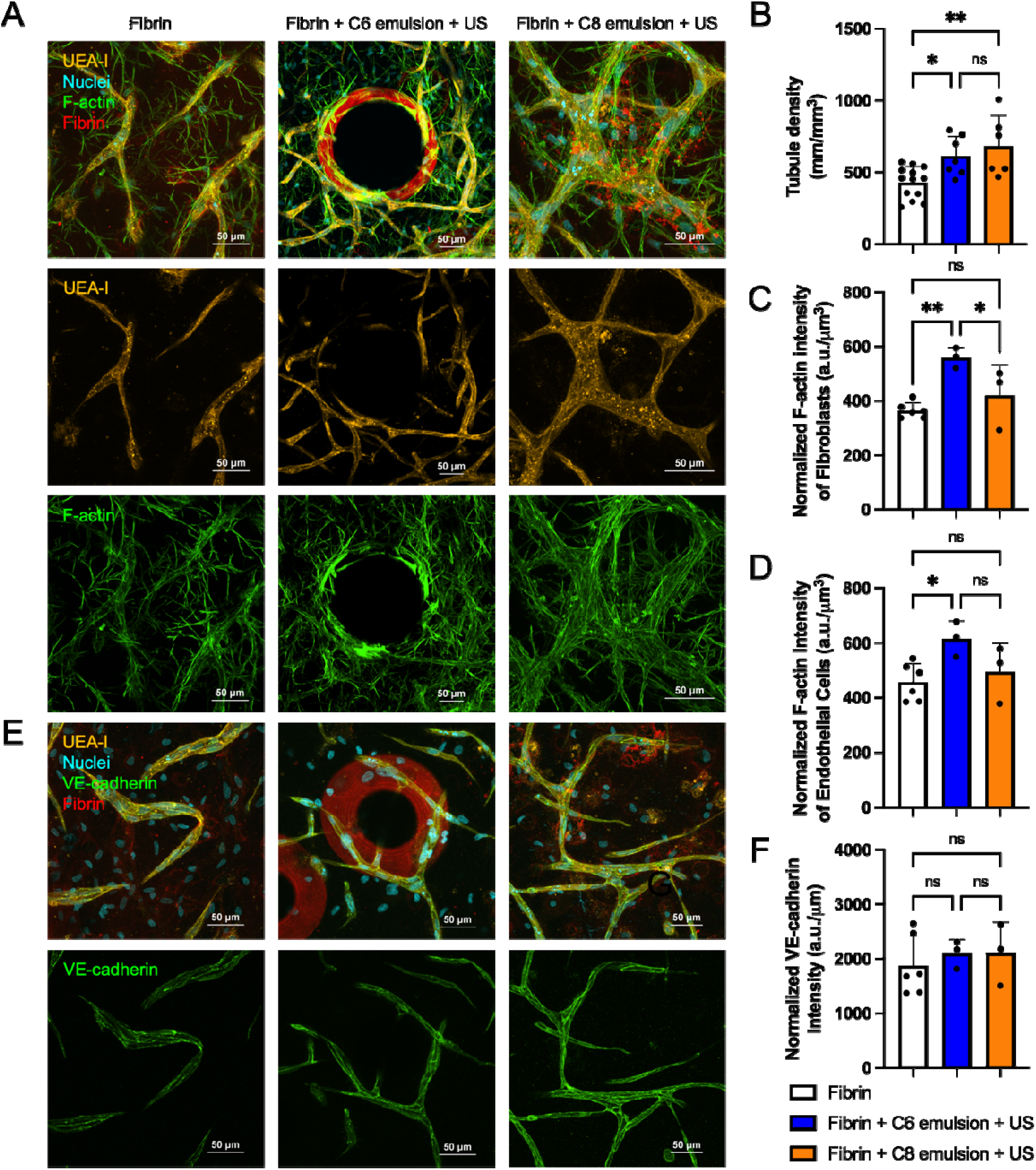
Ultrasound-driven matrix remodeling enhances endothelial sprouting and cytoskeletal organization in ARSs under full growth conditions. (A) Confocal images reveal endothelial sprouts in fibrin-only gels as well as in C6- and C8-ARSs containing ultrasound-generated bubbles and macropores, respectively, after 7 days of culture in growth media. In this study, emulsions did not contain bFGF. (B) Tubule density was quantified based on UEA-I staining. The intensity of F-actin in NHDFs (C) and HUVECs (D) was quantified based on phalloidin staining. (E) Confocal images also show VE-cadherin staining within the constructs. (F) The intensity of VE-cadherin was quantified based on the confocal images. N=3-13 gels per group with 5-10 fields of view per gel. Significant differences are denoted as follows: *p<0.05 and **p<0.01; ns denotes a non-significant difference. Additional images are found in Figure S5.

### 3.6. Morphological assessment of in vivo ARSs

Subcutaneously implanted ARSs and fibrin-only gels containing HUVECs and NHDFs were monitored longitudinally with B-mode ultrasound (**Fig. 7A, S6**). Across all groups, implant height decreased over time (**Fig. 7B**), with an average decrease ranging from 26-34% on day 1 to 38-54% on day 7. The echogenicity of ultrasound-treated C6-ARSs increased over time, with significantly greater differences seen on days 3 and 7 compared to day 0 (i.e., before ultrasound) (**Fig. 7C**). Additionally, similar differences were observed when comparing sonicated versus non-sonicated C6-ARSs on days 3 and 7. The echogenicity of other constructs remained stable throughout the study.

**Fig. 7.**
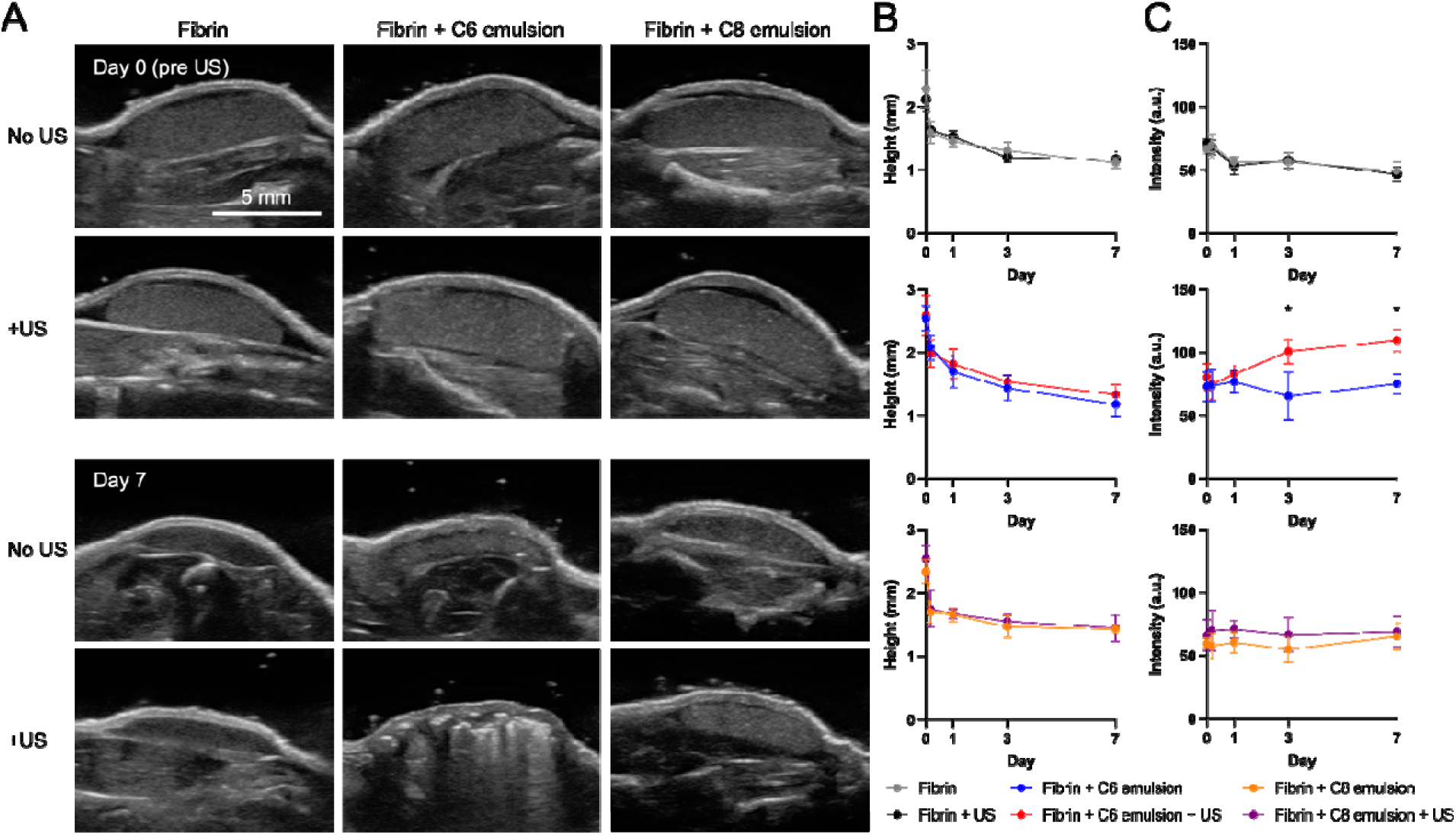
B-mode ultrasound imaging reveals in vivo compaction of the constructs. (A) B-mode ultrasound images of subcutaneously implanted, cell-loaded constructs on day 0 (i.e., immediately after implantation) and day 7. (B) Implant height and (C) intensity were longitudinally quantified over the 7-day study. N=4-5 implants per group with 3-5 fields of view per implant. Significant differences are denoted as follows: no US vs. +US (*p<0.05).

These B-mode findings were consistent with the appearance of the explants after harvest on day 7. H&E-stained images show bubble-containing regions in the sonicated C6-ARSs, whereas pore-like structures in the C8-ARSs were less obvious (**Fig. S7A**). Image analysis indicated that the average bubble and macropore diameters were approximately 150 µm and 11 µm, respectively (**Fig. S7B**). Together, these data show that ultrasound generated detectable in vivo remodeling of the fibrin ARS and that the resulting structural changes were preserved through the 7-day implantation period.

### 3.7. ADV-triggered release of bFGF stimulated vasculogenic assembly in vivo

Representative images show UEA-I+ structures in implants harvested on day 7, with more prominent endothelial signal in the ultrasound-treated C6- and C8-ARSs than in the corresponding untreated controls (**Fig. 8A, 8B**). Quantitatively, the UEA-I+ area increased significantly with ultrasound treatment in both the C6- and C8-containing groups, whereas the fibrin-only control did not show a significant change. However, there were no significant differences with quantified levels of Ter-119 within the implant (**Fig. S8**), an indicator of mouse erythroid cells, or the density of CD31+ vasculature in the granulation tissue surrounding the implant (**Fig. S9**). Additionally, no significant differences were observed in host cell migration based on nucleolin staining (**Fig. S10**). Collectively, these data suggest that ADV-mediated bFGF release was sufficient to enhance vasculogenic assembly by day 7, but did not impact angiogenesis surrounding the implant, anastomosis with host vasculature, or host cell migration.

**Fig. 8.**
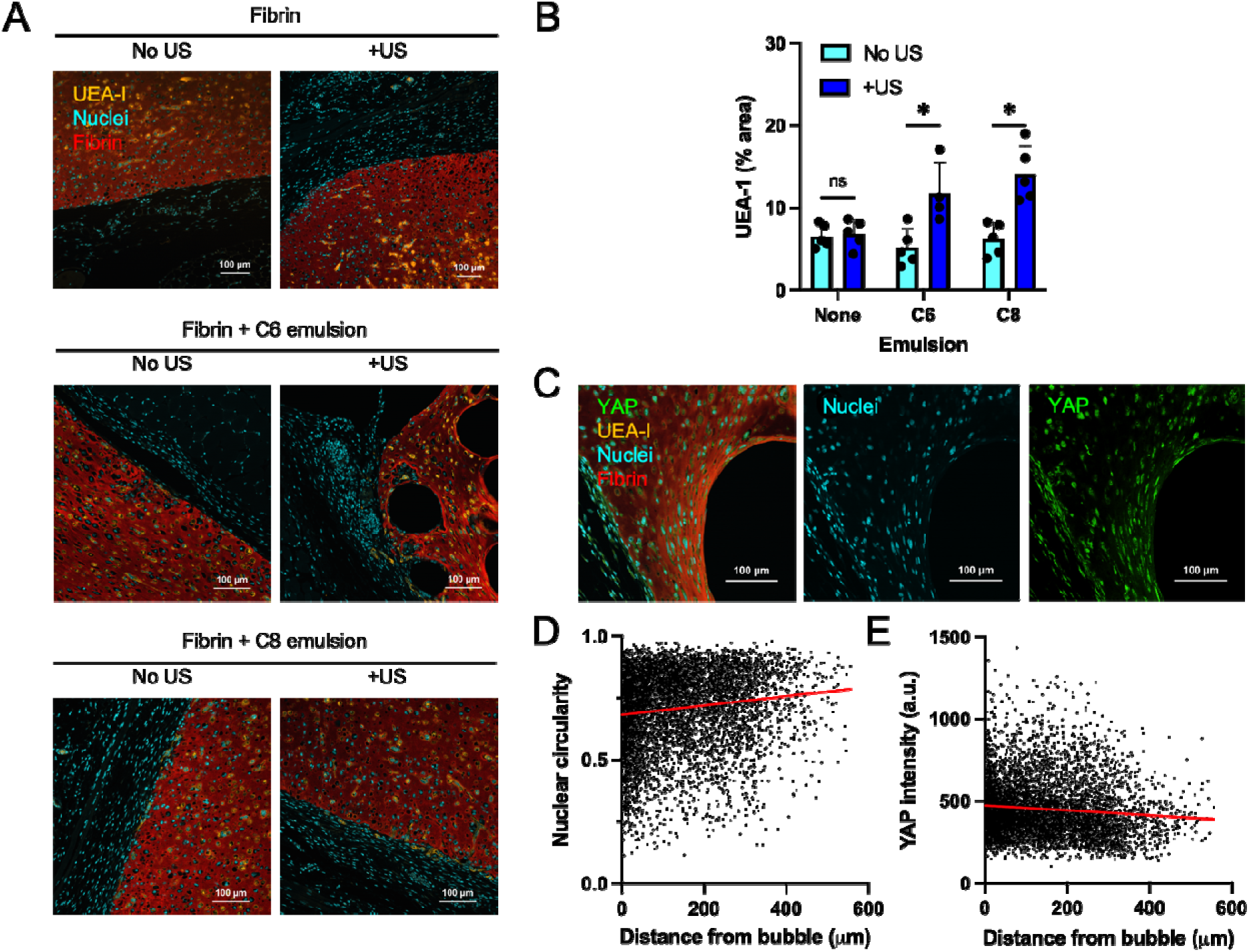
Ultrasound increased endothelial sprouting within ARSs while also impacting nuclear circularity and YAP levels. (A) In vivo cell-loaded constructs were explanted after 7 days, stained with UEA-I, and imaged with a confocal microscope to visualize endothelial structures. (B) Quantification of the UEA-I stained images revealed an increase in the area coverage of the endothelial structures within ARSs exposed to ultrasound (US). N=4-5 implants per group with 5-10 fields of view per implant. Significant differences are denoted as follows: *p<0.05; ns denotes a non-significant difference. (C) C6-ARSs exposed to US were immunostained for YAP. The confocal images show a region adjacent to an US-generated bubble. (D) Nuclear circularity and (E) YAP intensity exhibited positive and negative correlations, respectively, with distance from the bubble interface. The linear trends, shown by the red lines, were significant for both (p < 0.0001). N=5866 cells from 4 implants.

### 3.8. ADV-generated bubbles induced changes in nuclear morphology and YAP intensity

Given the dramatic restructuring induced by ADV-generated bubbles, we investigated nuclear morphology and YAP signaling within the harvested implants (**Fig. 8C**). Quantification of YAP intensity as a function of distance from each cell to the nearest bubble showed a significantly negative linear trend (p<0.0001) (**Fig. 8D**), indicating higher YAP signal in the region proximal to the compacted bubble interface. In parallel, nuclear circularity increased with distance from the bubble (p < 0.0001) (**Fig. 8E**), consistent with more elongated nuclei proximal to the bubble. These spatial trends indicate that bubble-induced, matrix restructuring produced localized changes to nuclear shape and YAP-related mechanotransduction.

## 4. Discussion

Hydrogels recapitulate the function of the extracellular matrix, thus providing critical biochemical and biophysical cues that impact vessel formation. Recent reviews highlight the design principles of hydrogels for vascular tissue engineering [30, 31]. Here, we developed an ultrasound-responsive hydrogel that enables simultaneous control of growth factor release and restructuring of the fibrin matrix. The phase-shift double emulsion within the ARS serves as both the growth factor depot and modulator of fibrin architecture. Ultrasound robustly increased bFGF release from the ARS compared with non□ultrasound controls, with the temporal profile strongly dependent on the perfluorocarbon species within the emulsion formulation (**Fig. 1D, 1E**). C6 emulsions produced an early burst followed by a rapid decay, while C8 emulsions supported more sustained and re□stimulatable release with repeated insonation. These findings are consistent with prior work demonstrating that ADV-mediated release of bFGF from acellular ARSs promoted endothelial network formation and angiogenesis [22, 32, 33].

Relative to conventional fibrin-based delivery systems - where bFGF is either mixed directly in the gel precursor prior to polymerization or immobilized via heparin or covalent binding - an ARS allows on-demand, temporally gated release. Conventional fibrin□bFGF constructs generally show monotonic release governed by diffusion and fibrin degradation, which can lead to an initial burst followed by subtherapeutic levels [34, 35]. The current platform therefore offers a degree of temporal programmability that is difficult to achieve with static formulations, and matches or exceeds the efficacy of vascular regeneration reported for other bFGF delivery vehicles in recent biomaterials studies [36].

Mechanistically, the difference between C6- and C8-emulsion behavior reflects their distinct boiling points, gas solubilities, and bubble stability. C6 (bulk boiling point of ∼56°C) generates bubbles that are stable following cessation of ultrasound and remain trapped within the fibrin matrix (**Fig. 2A, 2B**). Diffusion of dissolved gases from the surrounding microenvironment into the generated bubble [37] as well as the high solubility of oxygen within perfluorocarbon liquids [38] assist with bubble stability. Bubble growth leads to radial compaction of the surrounding fibrin matrix that ultimately stiffens the ARS at a bulk level (**Fig. 2E**). Characterization of bubble-induced stiffening via atomic force microscopy reveal hyperlocal changes to the modulus, corresponding to the bright (i.e., compacted) matrix around the bubble [39, 40]. Unlike prior studies that utilized higher concentrations of C6 emulsion within acellular gels (e.g., 0.5-1.0% (v/v)), these studied utilized 0.02% (v/v). Lower concentrations of C6 emulsion yield more rapid release kinetics than higher concentrations [29]. The bubble density within C6-ARSs post ultrasound exposure was sufficiently high to create an increase in pixel brightness within the implant and acoustic shadowing in vivo (**Fig. 7B, S6**), thus further highlighting why multiple ultrasound exposures were not used to trigger additional bFGF release from the C6-ARS. Despite these acoustic effects, the bubble density was insufficient to hinder bulk matrix compaction as observed in ARSs with 1% (v/v) emulsion [41].

Bubbles generated from C8-emulsion also induced some localized strain stiffening but ultimately recondensed following ultrasound exposure, due to the elevated bulk boiling point of C8 (i.e., 100°C). This led to the production of macropores with bright peripheries within the fibrin matrix (**Fig. 3A**). The partial release of payload from the C8-emulsion during ADV, coupled with the lack of stable bubble formation, enabled pulsatile bFGF release that could be reactivated with multiple ultrasound exposures [42] (**Fig. 1E**). Linked to drug release, repeated sonication generated macropores that grew, occupying a greater volume fraction within the ARS and merging adjacent macropores. Though high concentrations of porogens decrease bulk stiffness [43, 44], the concentration of ADV-generated macropores did not impact bulk rheological properties (**Fig. 3G, 3H**). Compared with other approaches to introduce macroporosity within hydrogels [45] - such as sacrificial templates, gas foaming, or bulk enzymatic degradation - an ARS offers two potential advantages. First, macropores can be generated on demand after implantation, avoiding potential issues with reduced construct handleability during implantation. Second, macropore size and density can be modulated by insonation parameters rather than fixed at the time of fabrication.

A critical result of this study was that ADV could be generated within ARSs without significantly impacting the viability of encapsulated HUVECs and NHDFs (**Fig. 4, S1, S2, S3**). Thus, observed differences in vasculogenesis were indicative of ultrasound-induced bFGF release and/or matrix restructuring rather than large-scale, ultrasound-induced cytoxicity. In all constructs, fibroblast viability was higher than endothelial viability at early timepoints (e.g., day 1, 4). This is consistent with other studies demonstrating the relative sensitivity of endothelial cells when initially encapsulated within hydrogels [46, 47]. There were small but significant reductions in viability observed in C8-ARSs exposed to two (i.e., days 0 and 4) and three (i.e., days 0, 2, and 4) ultrasound exposures. A potential hypothesis for this observation is that cells become more susceptible to ultrasound-induced deformation and associated cytotoxicity as the cells spread and interact with the fibrin matrix [48].

bFGF release from ARSs produced robust endothelial tubule formation in vitro (**Fig. 5C**) and in vivo (**Fig. 8B**), with the former yielding results in accordance with the dose response (**Fig. 5A**). In vitro tubule density was highest for C6-ARSs exposed to ultrasound and C8-ARSs with two and three exposures. Spatial gradients of bFGF are known to impact vascular formation [49]. In this study, there was a small but statistically significant correlation between tubule length and tubule-bubble distance (**Fig. 5D**). In a previous study, ADV in C6-ARSs led to incomplete disruption of the primary emulsion, thereby generating aggregates of W_1_ phase within the matrix surrounding the bubble [29]. Diffusion of bFGF from these aggregates, combined with the high binding affinity of bFGF for fibrin [50], could have contributed to the growth of longer tubules adjacent to the bubble. Localized effects of bFGF release were also observed when ADV was spatially patterned within subcutaneously implanted ARSs, producing directed angiogenic responses within the surrounding granulation tissue [22]. A key difference with this prior acellular study is that ultrasound-mediated bFGF release did not impact the density of blood vessels in the granulation tissue (**Fig. S9**). Thus, the effects of released bFGF were on the cells encapsulated within the ARSs.

Ultra-high-speed particle tracking studies in ARSs reveal that bubbles created from C6-and C8-emulsions generate radial strains of approximately 90% during ADV [51]. This is considerably higher than the 5-20% cyclic strain that is physiologically relevant for blood vessels and can change endothelial gene expression, signaling, and alignment [52, 53]. However, it is important to note that in this study, cells were exposed to 12 μs bursts of ultrasound at a rate of 100 Hz, which is equivalent to a 0.12% duty cycle. Following cessation of ultrasound, strain decreases dramatically. For C8-ARSs, the generated macropores do not longitudinally change in size [27], in the absence of cell-and enzyme-mediated remodeling. For C6-ARSs, bubble growth is slow. Based on the measured mean diameters (**Fig. 2C**) and an initial bubble diameter of 36.5 μm, which assumes a five-fold increase in particle size after ADV [54], bubble growth rates were 3.7, 2.6, and 1.4 nm/s across days 0 to 1, 1 to 4, and 4 to 7, respectively.

Without concomitant bFGF release, ultrasound-mediated bubble formation in C6-ARSs caused an increase in tubule density (**Fig. 6A, 6B**). This is consistent with bubbles creating a curved surface within the fibrin matrix that drives endothelial cell behaviors like migration [55]. However in addition to inducing curvature, bubbles also locally compact and stiffen the matrix, the latter of which has been shown to induce mixed effects on vascular formation [56, 57]. In parallel, F-actin levels increased for both HUVECs and NHDFs. Matrix stiffness and tension are established drivers of F-actin polymerization and stress fiber formation for both fibroblasts [58] and endothelial cells [59]. Tubule formation was also enhanced by the generation of macropores within C8-ARSs, thus supporting the hypothesis that macropores facilitate endothelial growth and migration [60]. Though, matrix densification was also observed at the periphery of macropores (**Fig. 3A**), the width of the compacted region was qualitatively less than in C6-ARSs (**Fig. 2A**), which may explain why increases in F-actin were not observed in C8-ARSs. Overall, ultrasound induces simultaneous changes to matrix architecture and mechanical properties within an ARS, which can both impact tubule formation.

Prior studies demonstrate that the integrity of VE-cadherin junctions was weakened by increasing substrate stiffness [61] and heterogeneity in matrix stiffness [62]. Yet, significant differences in VE-cadherin intensity were not observed in any of the experimental conditions (**Fig.7F**). VE-cadherin, the principal adhesion protein at endothelial adherens junctions, is required for vascular integrity and for regulating permeability during both developmental and tumor angiogenesis [63, 64]. During sprouting, its function is mainly controlled by phosphorylation, endocytosis, and local turnover at junctions, rather than by large changes in total protein level. Studies show that VEGF, a component of the growth media used in these studies, promotes endothelial junction remodeling by driving actin-dependent redistribution of VE-cadherin to new adhesion sites rather than increasing its overall expression, enabling dynamic cell elongation and sprouting [65, 66]. In addition, VE-cadherin forms mechanosensory complexes with PECAM-1 and VEGFR2 that regulate junctional tension and turnover in response to mechanical forces while maintaining relatively stable total VE-cadherin levels [67]. Overall, this suggests that ultrasound-induced matrix restructuring enhanced tubule density and F-actin levels without impacting VE-cadherin.

In addition to stiffening the fibrin matrix, ADV-generated bubbles modulated YAP signaling and nuclear shape, with negative and positive correlations as a function of cell-bubble distance, respectively (**Fig. 8D, 8E**). The opposing correlations are consistent with prior studies demonstrating that nuclear elongation led to increased YAP translocation [68]. Additionally, these patterns align with studies showing that increased matrix stiffness and cytoskeletal tension promote YAP/TAZ nuclear localization, drive endothelial sprouting, and regulate vascular remodeling via integrin□ and thrombospondin□1□mediated mechanotransduction [19, 69, 70]. In particular, endothelial mechanobiology studies using vessel□on□chip models have demonstrated that stiff matrices impede shear□induced YAP responses but enhance stiffness□driven nuclear YAP accumulation and angiogenic signaling [71]. Our findings are compatible with this paradigm: ADV□generated zones of matrix compaction likely increased local stiffness and tension, promoting nuclear mechanotransduction even in the absence of shear, and thereby supporting longer tubules and sprouting around the bubble interface (**Fig. 5D**). This provides a plausible mechanistic link between ultrasound□induced structural remodeling and the observed vasculogenic enhancement, complementing the bFGF delivery mechanism.

There are some limitations of this study that warrant consideration. First, in vivo experiments were conducted with subcutaneous implants and harvested after 7 days. The subcutaneous model does not mimic the hypoxic environment found within ischemic tissue. Additionally, the 7-day endpoint precluded the study of host anastomosis (**Fig. S8**). Without prevascularization, connection to host vessels can take two weeks for fibrin hydrogels containing endothelial cells and support cells like fibroblasts [72] or mesenchymal stromal cells [56]. Second, ADV-mediated drug release and matrix remodeling are inherently linked, thereby preventing disentanglement of biochemical and biophysical cues on vasculogenic assembly within the ARS. Future studies should investigate long-term vessel stability and evaluation within more physiologically relevant disease models (e.g., hind limb ischemia model). Extending the platform to more complex tissues (e.g., cardiac or skeletal muscle) and scaling to larger volumes will also be important for translational impact.

## 5. Conclusions

We developed an ultrasound-responsive, fibrin-based, composite hydrogel that supported vasculogenic assembly. The phase-shift double emulsion within the ARS enabled ultrasound-controlled release of bFGF in combination with changes to matrix architecture and mechanical properties. bFGF release kinetics and matrix restructuring were highly dependent on the perfluorocarbon species within the emulsion. Upon a single ultrasound exposure, bFGF was released from C6-ARSs, yielding stable bubbles that locally compacted the matrix and increased macroscale stiffness. Conversely, macropores were generated in C8-ARSs, which did not impact macroscale viscoelasticity, and enabled repeated release of bFGF using multiple ultrasound exposures. ADV could be generated in ARSs without significantly impacting the viability of 3D encapsulated HUVECs or fibroblasts. Ultrasound-mediated release of bFGF increased vasculogenic assembly in both in vitro and in vivo models. In the absence of bFGF release, ADV also increased vasculogenic assembly in C6- and C8-ARSs, with changes in F-actin and YAP noted for C6-ARSs. Overall, these results highlight the potential for ARSs to actively shape vascular microenvironments in regenerative medicine.

## 6. Acknowledgements

This work was supported by NIH grant R01HL139656.

