## Supplemental for "Ultrasound-reconfigurable scaffolds enable dynamic control of biochemical and biophysical cues for vascular network formation"

2026-08-13

^1^Department of Radiology, University of Michigan, Ann Arbor, MI, USA

^2^Department of Biomedical Engineering, University of Michigan, Ann Arbor, MI, USA

^3^Applied Physics Program, University of Michigan, Ann Arbor, MI, USA

*Corresponding Author

Mario L. Fabiilli

University of Michigan

1301 Catherine Street

6436C Medical Sciences Building I

Ann Arbor, MI, 48109-5667, USA

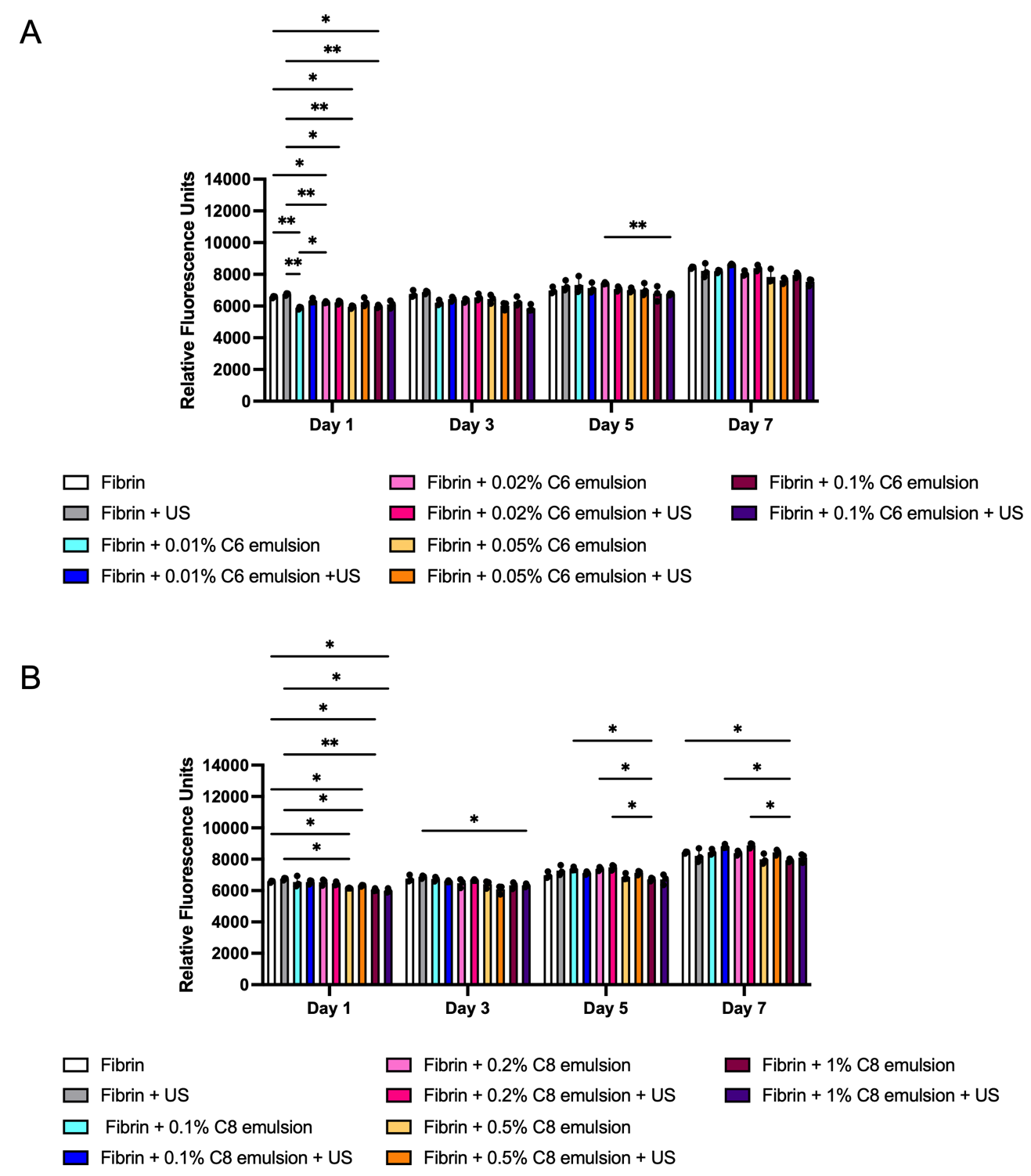

**Fig. S1. Metabolic activity of NHDF monoculture within ARSs of varying emulsion concentrations.** NHDFs were encapsulated within ARSs containing (A) C6- or (B) C8-emulsions. The Alamar blue assay was used to longitudinally evaluate metabolic activity as a function of emulsion concentration and ultrasound (US) exposure. For C6-emulsion, the highest average metabolic activities were observed at 0.01% and 0.02% (v/v), especially when ultrasound was applied. For C8-emulsion, the highest activities were observed at 0.1% and 0.2% (v/v). This difference in concentration between C6 and C8 emulsions is linked to their post-US morphologies (i.e., bubbles versus macropores). N=3 gels per group. Significant differences are denoted as follows: *p<0.05 and **p<0.01.

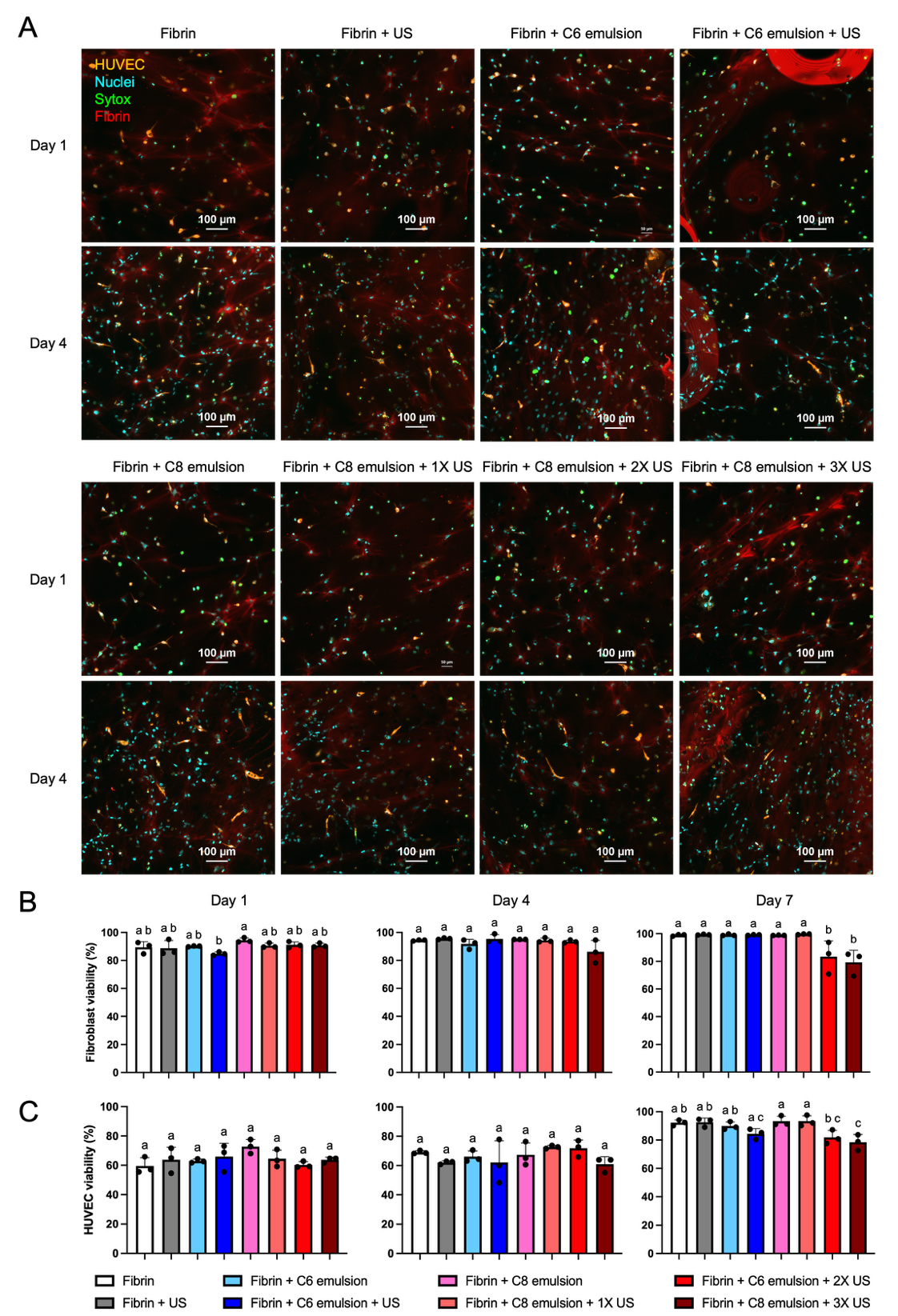

**Fig. S2. Co-culture viability in ARSs.** (A) Confocal images show RFP-HUVECs and NHDFs co-encapsulated within fibrin-only gels as well as ARSs with C6- or C8-emulsion after 1 and 4 days of culture. Viability was assessed using Sytox-staining. The viability of NHDFs (B) and HUVECs (C) were quantified. A custom script was used to classify the cells as follows: live HUVEC (+RFP, -Sytox), dead HUVEC (+RFP, +Sytox), live NHDF (-RFP, -Sytox), and dead NHDF (-RFP, +Sytox). N=3 gels per group with 5-10 fields of view per gel. Statistically significant differences in panels B and C are denoted in compact letter display.

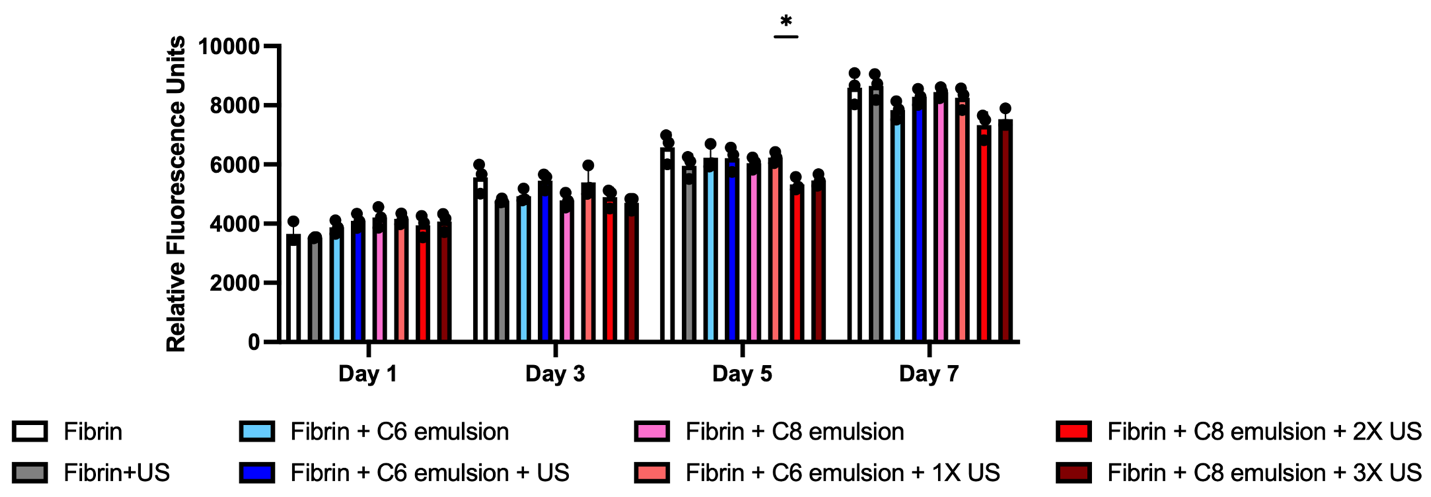

**Fig. S3.** **Metabolic activity of HUVEC-NHDF co-culture within ARSs following ultrasound exposure.** Alamar blue assay was used to quantify metabolic activity in C6- and C8-ARSs containing 0.02% and 0.2% (v/v) emulsion, respectively. C6-ARSs were exposed to ultrasound (US) on day 0. C8-ARSs were exposed to US on day 0 (1X US), days 0 and 4 (2X US), and days 0, 2, and 4 (3X US). N=3 gels per group. Significant differences are denoted as follows: *p<0.05.

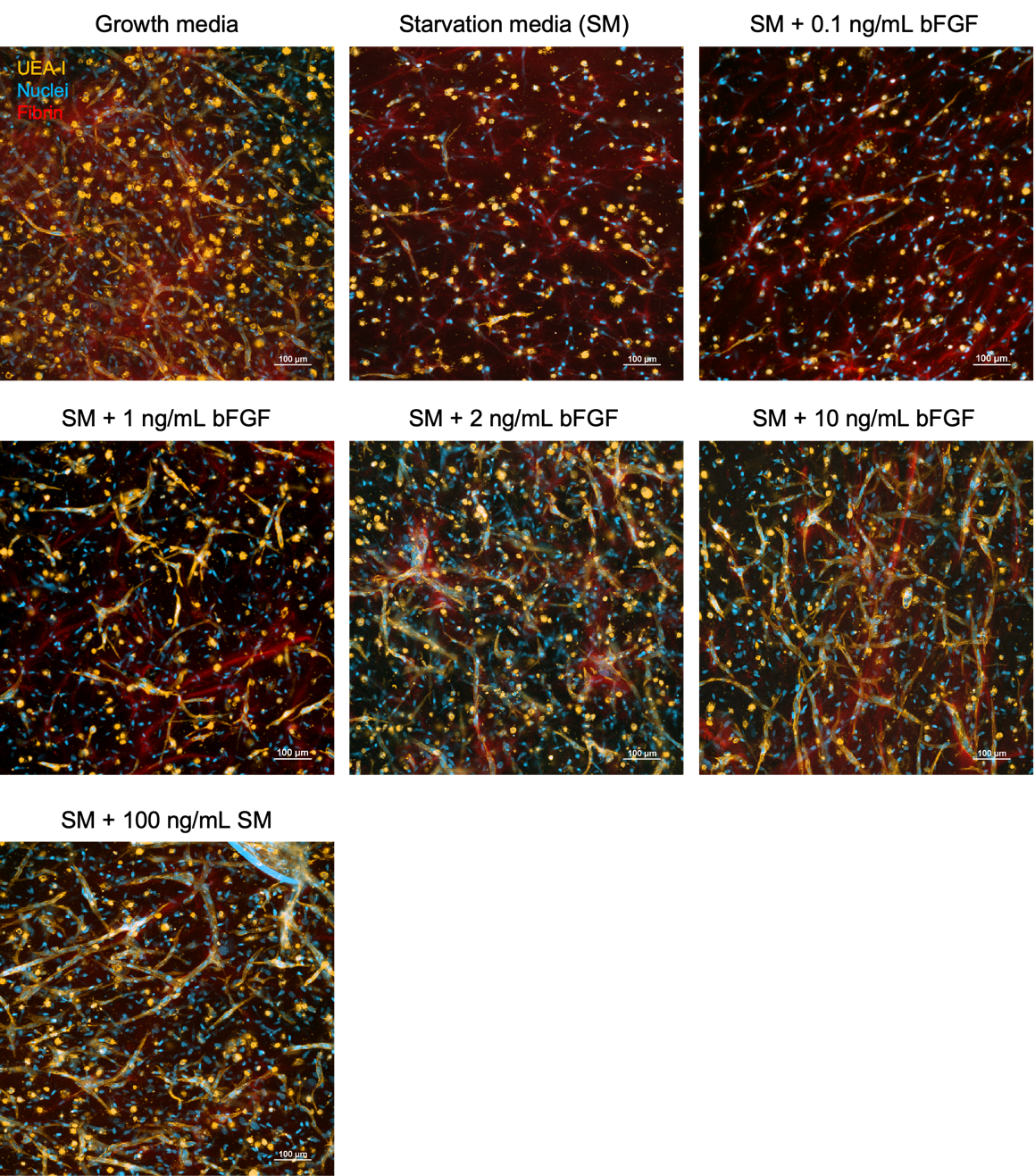

**Fig. S4. Endothelial sprouting was dependent on bFGF concentration.** HUVECs and NHDFs were encapsulated within fibrin-only hydrogels at 5 x 10^5^ cells per mL for each cell type. Constructs were cultured for 7 days in either fully supplemented growth media, starvation media, or starvation media supplemented with varying concentrations of bFGF. Constructs were subsequently stained with UEA-I and imaged with a confocal microscope to evaluate endothelial sprouting and tubule formation.

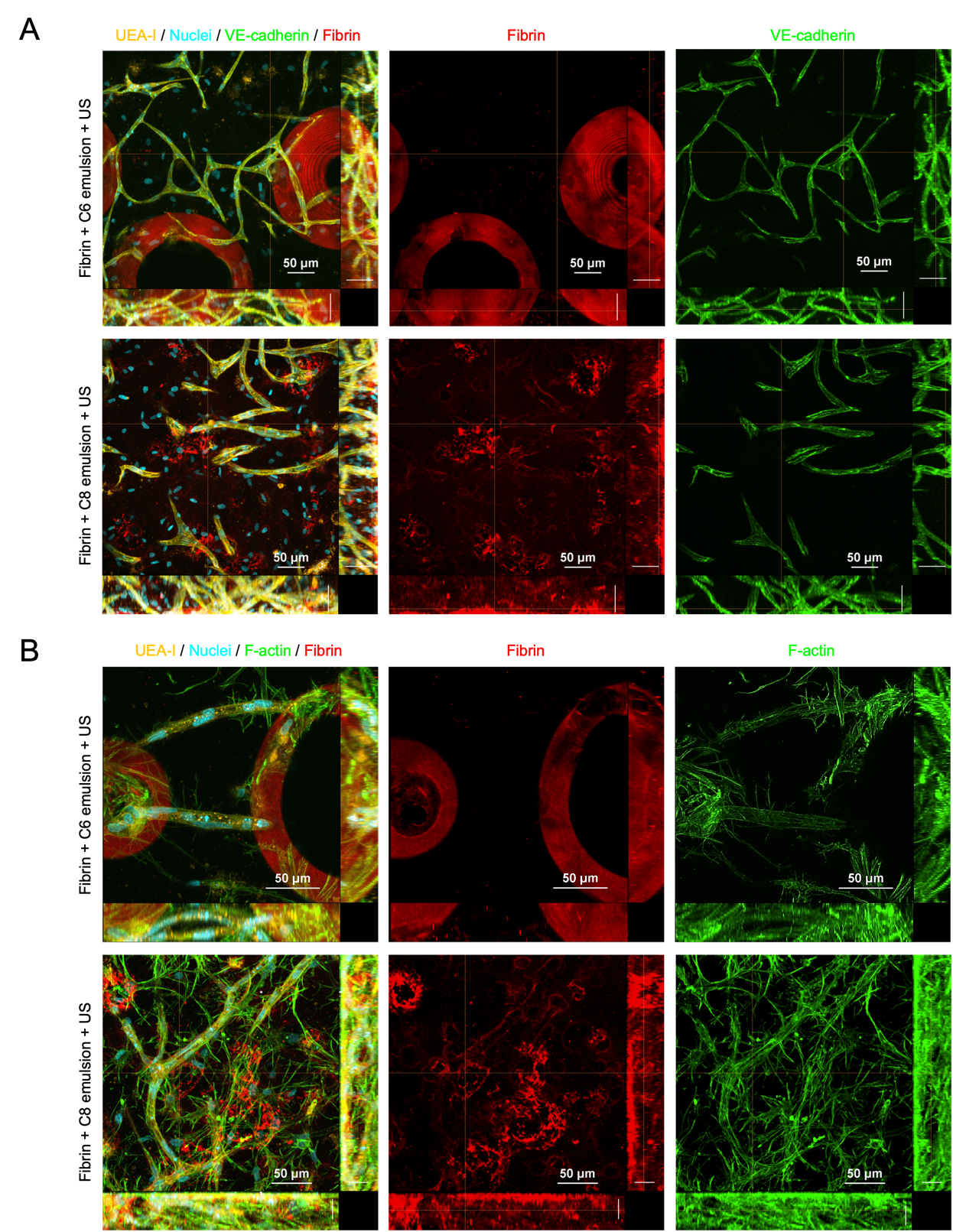

**Fig. S5. Confocal images show the morphology of endothelial tubules adjacent to ultrasound-generated bubbles and macropores within C6- and C8-ARSs, respectively.** Cell-laden ARSs were stained for VE-cadherin (A) and (B) F-actin. Within each subpanel, the larger image in the top left is a maximum intensity project image. The rectangular images on the bottom and right sides correspond to images within the XZ and YZ planes. The vertical and horizontal lines within the large image denote the location of the YZ and XZ planes, respectively. These images support the observation that ADV-generated bubbles and macropores create localized zones of matrix remodeling associated with enhanced F-actin organization and endothelial tubule sprouting.

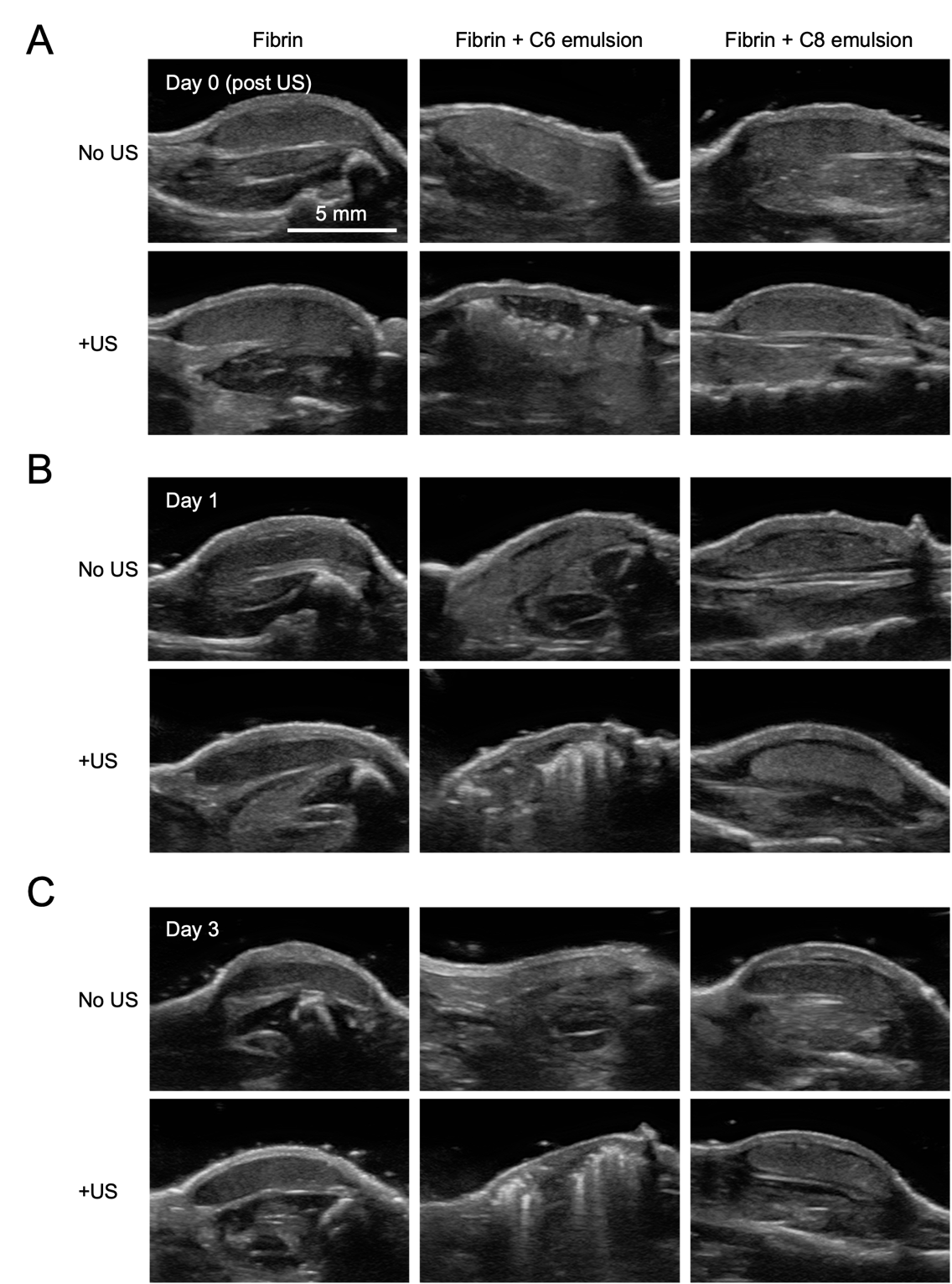

**Fig. S6. B-mode ultrasound images of subcutaneously implanted fibrin-only constructs and ARSs.** Images from (A) day 0 (post ultrasound), (B) day 1, and (C) day 3 are shown. All constructs include HUVECs and NHDFs. Note the increase in echogenicity (i.e., brightness) within the C6-ARS exposed to ultrasound (US) following bubble formation.

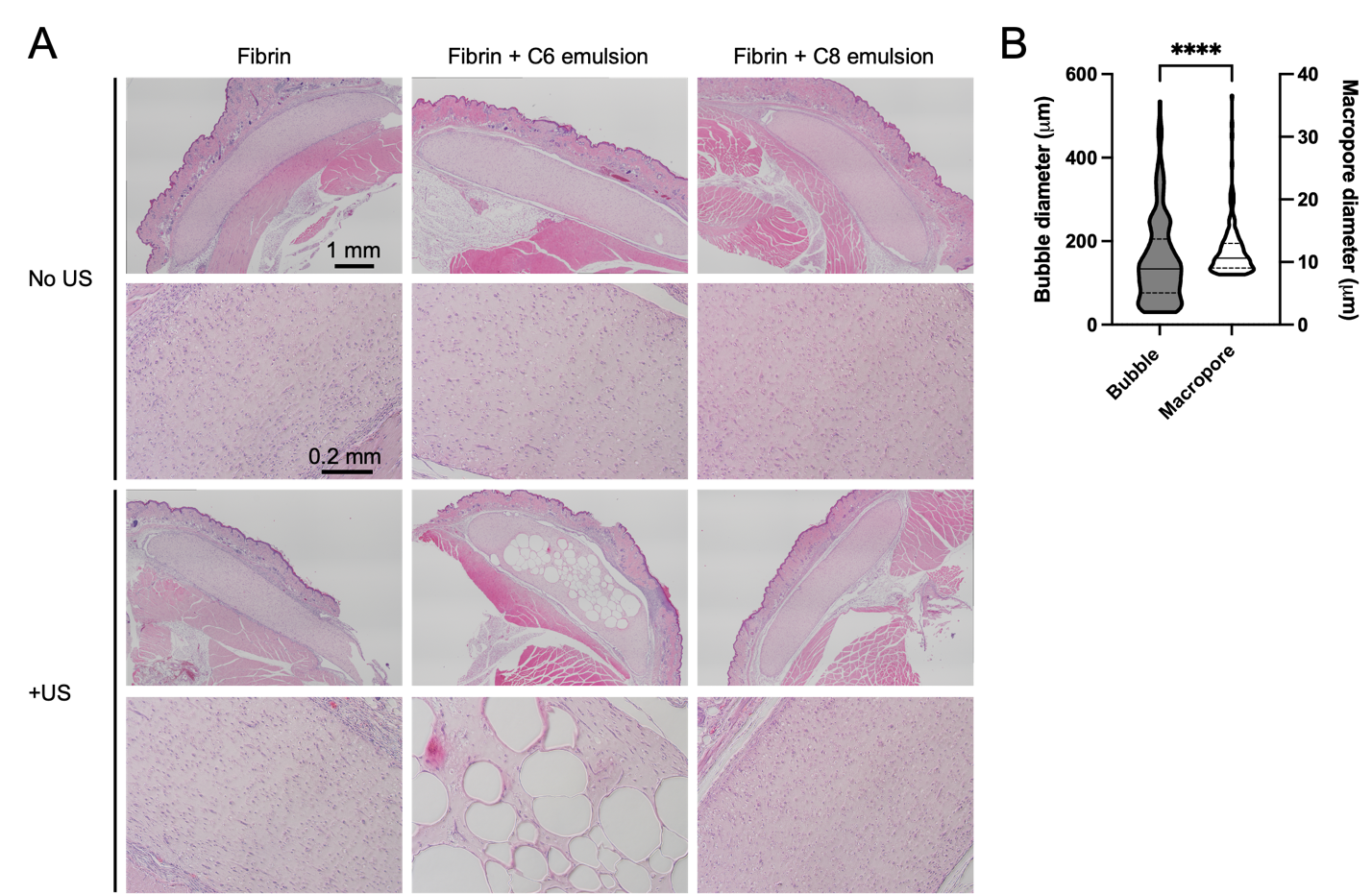

**Fig. S7. Histological assessment of in vivo bubble and macropore formation in ARSs.** (A) In vivo constructs were explanted after 7 days and stained with hematoxylin and eosin (H&E). Bubbles are clearly visible in the C6-ARSs exposed to ultrasound (US). (B) The diameters of ultrasound-generated bubbles and macropores were quantified. Data are shown as violin plots with the first quartile, median, and third quartile denoted. N=4-5 implants per group with 5-10 fields of view per implant. Statistically significant differences are denoted as follows: ****p<0.0001.

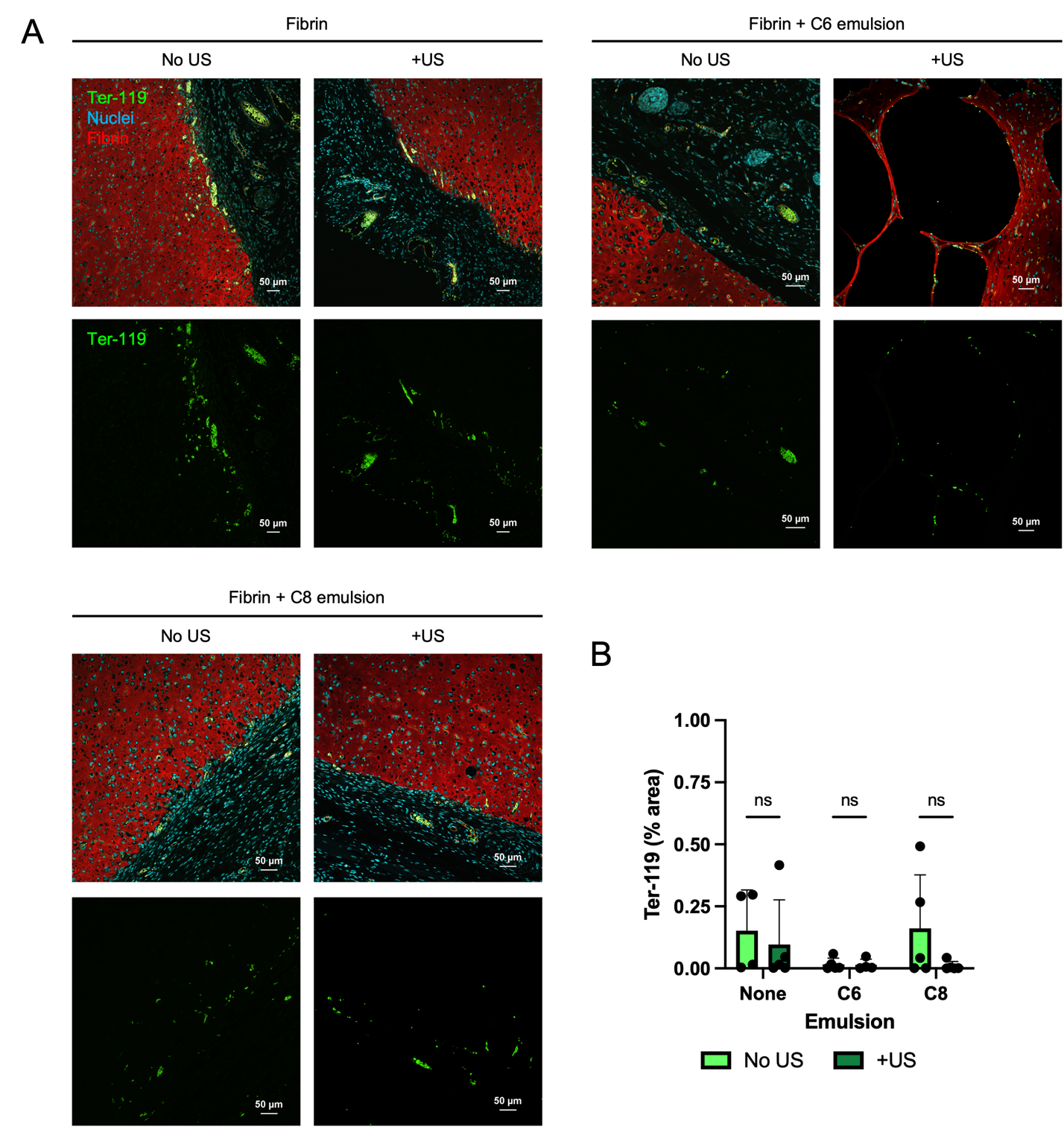

**Fig. S8. Immunostaining of mouse erythroid cells within the implant.** (A) In vivo constructs were explanted after 7 days and immunostained for Ter-119, a marker of mouse erythroid cells (B) The area of Ter119+ staining within the implant was quantified and normalized by the total implant area. N=4-5 implants per group with 5-10 fields of view per implant. Ns denotes a non-significant difference.

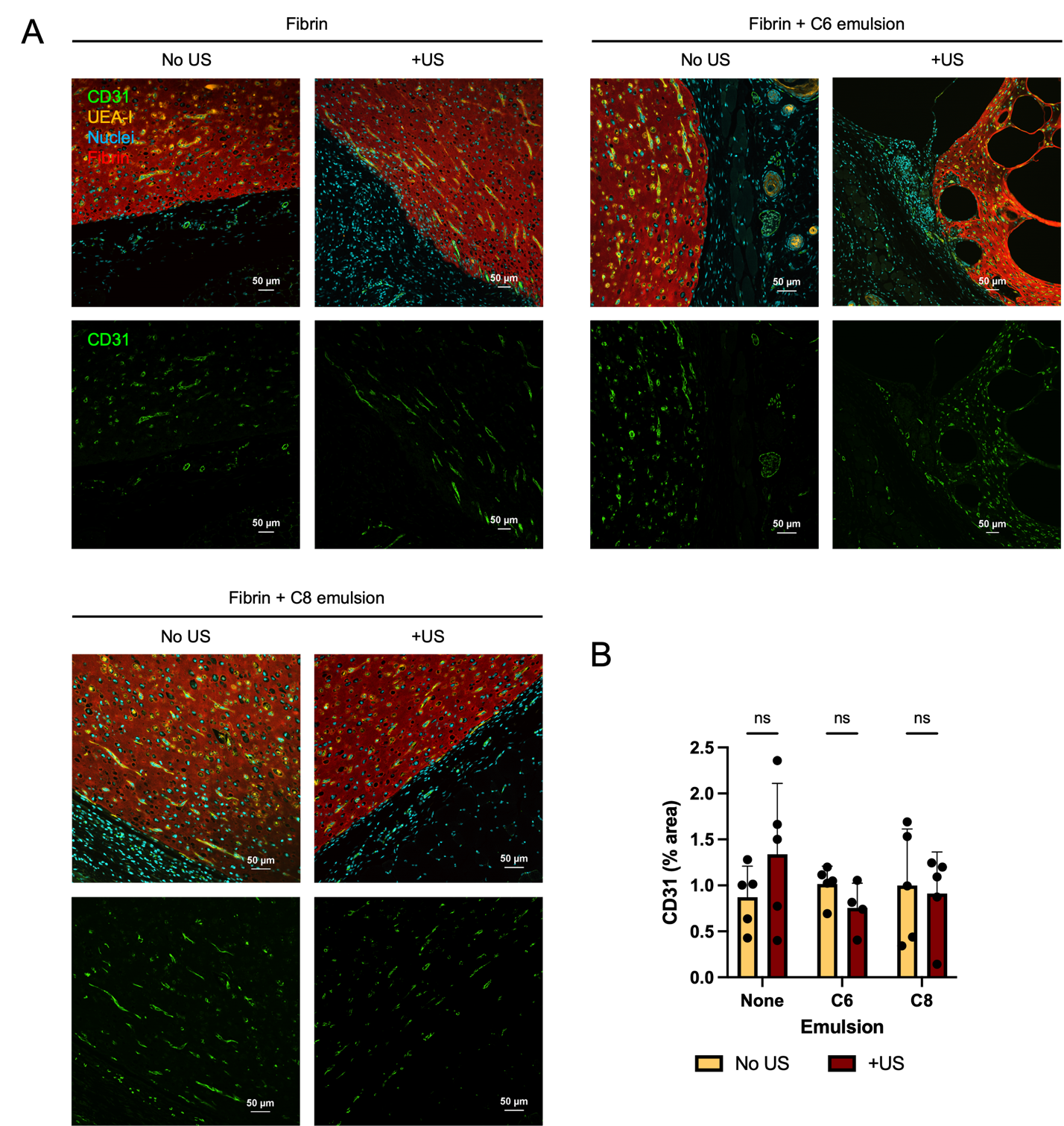

**Fig. S9. Immunostaining of vascularization around the implant.** (A) In vivo constructs were explanted after 7 days and immunostained for CD31, which stained both mouse- and human-derived blood vessels. (B) The area of CD31+ staining within the granulation tissue surrounding the implants was quantified. The upper region of granulation tissue was defined as the region of tissue between the panniculus carnosus and upper implant boundary. The lower region of granulation tissue was defined as the region of tissue between the lower implant boundary and skeletal muscle. Upper and lower regions were aggregated and used to normalize the CD31+ area. N=4-5 implants per group with 5-10 fields of view per implant. Ns denotes a non-significant difference.

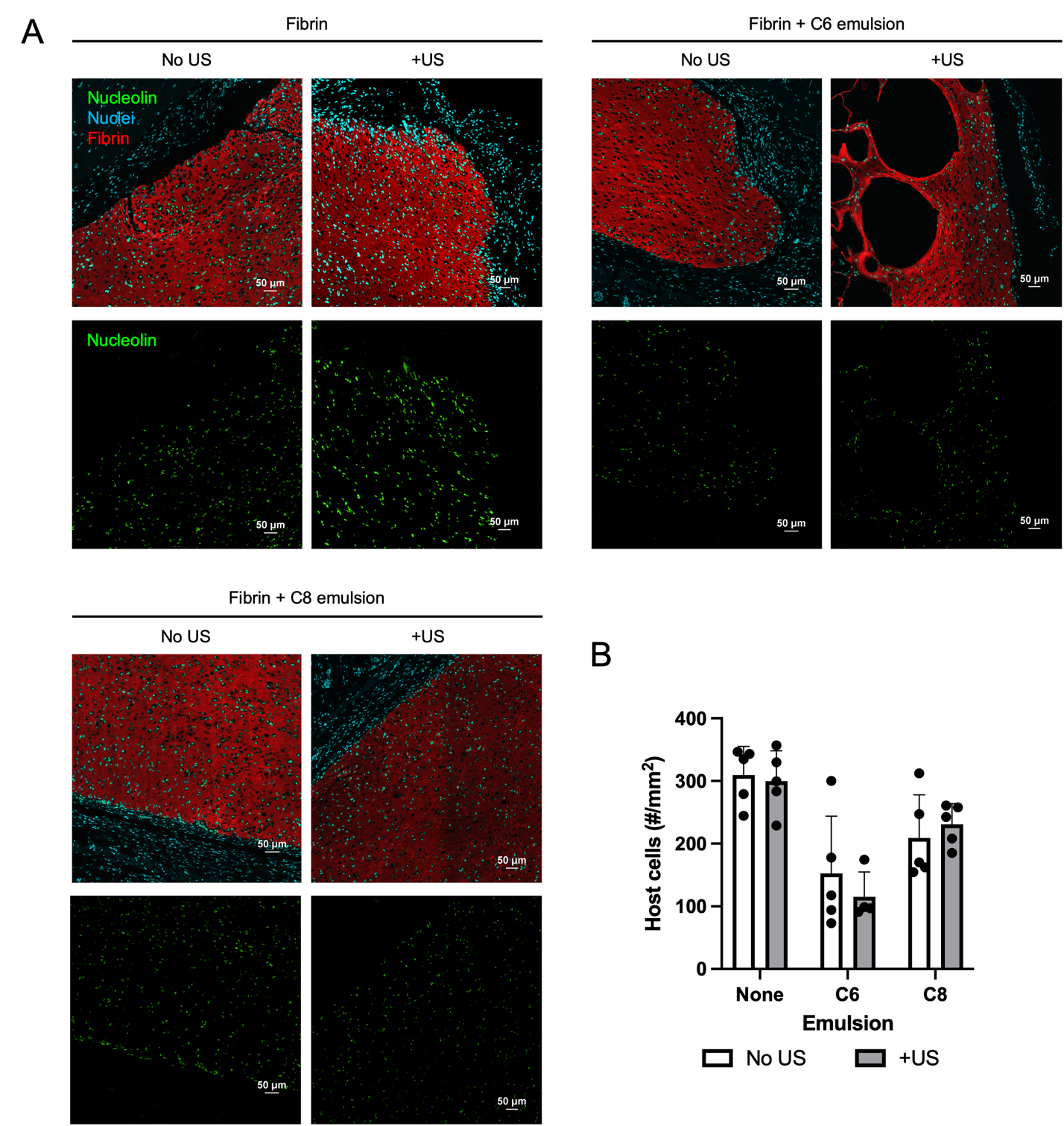

**Fig. S10. Immunostaining of host cell invasion.** (A) In vivo constructs were explanted after 7 days and immunostained for human nucleolin. (B) Host cell migration was quantified based on the nucleolin-stained images. The number of host cells was calculated as the difference between the number of DAPI+ cells and nucleolin+ cells within the implant. Host cell number was normalized based on the implant area. N=4-5 implants per group with 5-10 fields of view per implant.
